# Shaping the Genome via Lengthwise Compaction, Phase Separation, and Lamina Adhesion

**DOI:** 10.1101/2022.02.28.482402

**Authors:** Sumitabha Brahmachari, Vinícius Contessoto, Michele Di Pierro, José N. Onuchic

## Abstract

The link between genomic structure and biological function is yet to be consolidated, it is, however, clear that physical manipulation of the genome, driven by the activity of a variety of proteins, is a crucial step. To understand the consequences of the physical forces underlying genome organization, we build a coarse-grained polymer model of the genome, featuring three fundamentally distinct classes of interactions: lengthwise compaction, i.e., compaction of chromosomes along its contour, self-adhesion among epigenetically similar genomic segments, and adhesion of chromosome segments to the nuclear envelope or lamina. We postulate that these three types of interactions sufficiently represent the concerted action of the different proteins organizing the genome architecture and show that an interplay among these interactions can recapitulate the architectural variants observed across the tree of life. The model elucidates how an interplay of forces arising from the three classes of genomic interactions can drive drastic, yet predictable, changes in the global genome architecture, and makes testable predictions. We posit that precise control over these interactions in vivo is key to the regulation of genome architecture.

## I. INTRODUCTION

Chromosomes are long polymers, whose three-dimensional architecture is regulated by a myriad of proteins, including molecular motors. The architectural features are reflected in the characteristic ensembles of conformations observed through the many variants of DNA-DNA proximity ligation assays, such as Hi-C [1–5], and high-resolution microscopy techniques [6, 7]. These experiments show that, while none of the ensemble structures are identical, chromosome architecture specific to cell type and different cell-cycle phases share common features. In a recent study, surveying genome architecture in multiple organisms spanning the tree of life, we found four commonly observed, classifying characteristics: territorial chromosomes, clustered centromeres, clustered telomeres, and centromere-to-telomere axis or chromosomes with aligned arms [8]. A variety of studies, ranging from single-molecule to bulk *in vivo*, allude to the regulation of genome architecture as a complex network of interactions involving a gamut of proteins, like SMC complexes, architectural proteins, and chromatin remodelers [5, 9–14]. Naked DNA is known to exhibit equilibrium-polymer-like properties, interactions with proteins, while preserving the overall integrity of the polymer, stabilizes a structure via constraining or enhancing selective polymer degrees of freedom. We postulate that genome-structure characterization occurs via the regulation of three classes of degrees of freedom: looping, topology-independent segregation or clustering, and tethering to the nuclear envelope or lamina. The underlying interactions driving these structural modes are, respectively, lengthwise compaction, self-adhesion among chromatin, and adhesion of chromatin with the nuclear lamina.

Lengthwise compaction represents a thermodynamic force that folds the chromosomes along its contour [15, 16], which has also been referred to as the ideal chromosome potential in data-driven models of chromosome [17, 18]. Chromatin looping, a pervasive *in vivo* feature, is regulated at lengthscales ranging a few kilo-basepairs (kb) to mega-basepairs (Mb) [5, 11]. When chromatin is coarse-grained, the loops smaller than the coarse-graining lengthscale of a monomer are inconsequential to the structure, but the tendency to form loops larger than the monomer size leads to an average compaction force along the chromosome contour. This compaction force underlies lengthwise compaction that drives the looping degrees of freedom and controls the decay of the contact probability between chromosome segments as a function of their genomic distance. SMC complexes, thanks to their loop extrusion and stabilization activity, are prominent drivers of chromatin looping probability and control lengthwise compaction of chromosomes [9, 11, 13, 19–22]. Models of chromosome have realized lengthwise compaction in different ways. Stochastic simulations have demonstrated that loop extrusion may control the contact probability decay [23–25], a signature of lengthwise compaction. Complimentarily, steady-state dynamics of coarse-grained polymer simulations have shown that the contact probability scaling can be recapitulated using pairwise-interaction free energies (Ideal Chromosome potential) that decay with the contour distance between the interacting pair [17, 18]. Depending on the desired level of coarse-graining, one method may be more applicable than others, however, a model of chromosomes is incomplete without lengthwise compaction.

The second principle, self-adhesion among chromatin blocks, drives phase separation of the self-adhering segments into compartments. The characteristic plaid-patterns of interactions between non-neighboring loci, as observed in HiC-data, correspond to compartmental segregation [2, 26]. These experiments also argue that the compartments are correlated with epigenetic modifications of chromatin, i.e., heterochromatin (defined by post-translational histone modifications, like H3K9me2/3) and euchromatin (defined by modifications like H3K27ac) segments segregate into separate compartments. The self-adhesion among segments, arising from inter-nucleosome adhesion [27] and aggregation by cross-linking proteins, such as HP1 [28], is reminiscent of polymeric or colloidal liquids in marginally bad solvent. This has led to the block copolymer models of chromatin where euchromatin and heterochromatin blocks feature enhanced self-adhesion, and lead to respective compartmentalization [18, 29–32]. We recently found clustering of constitutive heterochromatin (centromeres and telomeres) as a classifying characteristic of genome architecture, and postulated phase separation as the driving mechanism [8]. Importantly, unlike lengthwise compaction that is restricted to intra-chromosome interactions, chromatin self-adhesion only depends on the epigenetic character and is a chromosome-topology-independent mechanism of genome organization.

The third class of interactions constitutes chromatin blocks, also called Lamina-Associated Domains (LADs) [33–35], interacting preferentially with the nuclear envelope. LADs are repressive environment, rich in heterochromatin-specific histone modifications, harboring lower gene density [33, 34]. Proteins, like lamin B1 and lamin A/C in eukaryotes [35, 36] and cec-4 in worms [37, 38], are known to tether heterochromatin to the nuclear envelope. Like self-adhesion, this interaction depends on the epigenetics of the chromosome segments, however, unlike self-adhesion, tethering with the envelope is capable of reorganizing the relative positioning of chromosomes from the center of the nucleus to the periphery [29, 39].

We develop a theoretical framework to understand the effects of the three above mentioned forces of genome organization, and investigate if the competition among these forces can recapitulate experimental observations, such as the species-wide architecture variants observed at the chromosomal lengthscales [8]. Unfolded chromosomes are represented in our model as a homopolymer or an array of connected monomers (see Supplementary Materials). The monomers represent coarse-grained chromatin domains, containing 20-50 kb DNA, that have emerged as organizational units of chromosomes [40, 41]. Lengthwise compaction is implemented as an interaction potential that favors contact between intra-chromosome loci pairs with an intensity that decays with increasing genomic separation between the interacting loci (Fig. 1A-B), similar in spirit to the Ideal Chromosome potential [17, 18, 42]. This potential transforms the homopolymer into a lengthwise-compacted polymer (LCP) and controls the steepness of contact probability decay along the genomic contour (Fig. 1C), mimicking SMC complex activity [11, 13, 21]. The LCP potential is composed of two terms: short-range and long-range compaction. The strength of lengthwise compaction between loci pairs that are less than a characteristic length (about 10 monomers or 200-500 kb) apart on chromosome contour is mainly controlled by the short range compaction, whereas, intrachain monomer-pair interactions beyond this characteristic length are only controlled by the long-range component. These two components depict the activity of SMC variants: Condensin I, Condensin II, and cohesin. Varying the two components of the LCP potential we capture variations in the activity of SMC complexes, which can arise either due to altered concentration or varied residency time of SMCs on DNA. Condensin II is known to establish long chromatin loops and controls the contact probability between intra-chromosome segments that are many hundreds of kb to Mb apart [8, 9, 11, 21, 43, 44], which we postulate controls the long-range lengthwise compaction of chromosomes. Condensin I is associated with shorter loops of less than 100 kb [9, 11, 43, 44], which corresponds to the short-range lengthwise compaction in our set up. Cohesin activity may vary widely depending on the genomic sequence, as it interacts with factors like architectural proteins [5]. However, given the typical loop size associated with Cohesin is a few hundred kb [45], we simply associate Cohesin with short range lengthwise compaction. Varying the short- and long-range components of lengthwise compaction, our model describes the various chromosome structural phenotypes expected upon hyperactivity or depletion of these proteins [13, 46– 48].

**FIG. 1.**
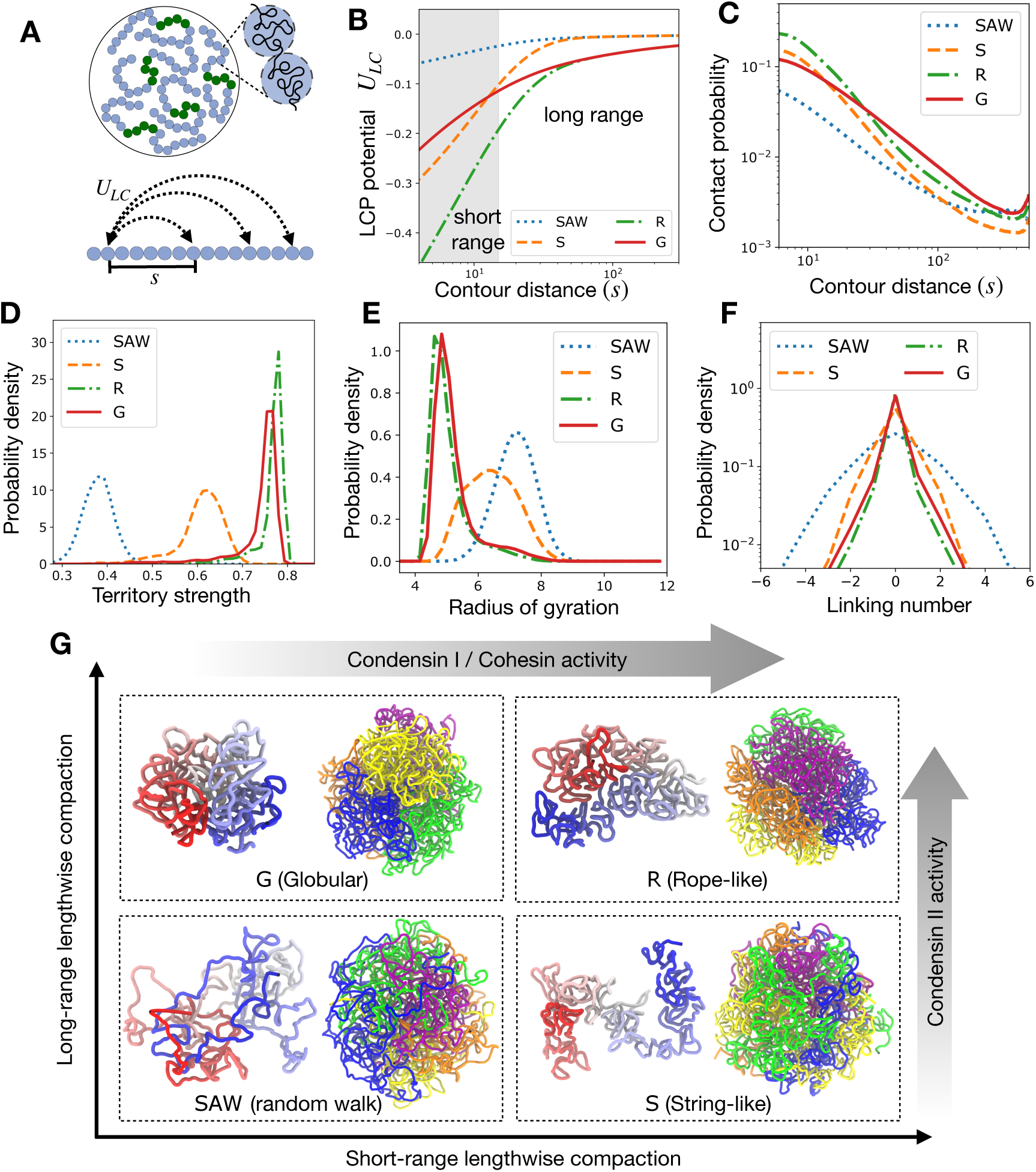
Regulation of chromosome structure and entanglements via lengthwise compaction. (A) Schematic of simulation set up. Five chromosomes, constituted of an array of 500 monomers each, are simulated, where the central regions shown in green are centromeres. (B) Lengthwise compaction potential *U*_*LC*_, plotted as a function of the genomic distance *s*. Note the generic decay with genomic distance, however, the intensity of the potential at long and short ranges are distinct for the different phenotypes: Random walk-like (SAW), Globular (G), Stringy (S), and Rope-like (R). (C) Probability of contact between loci pairs that are *s* distance apart along the contour, plotted as a function of *s*. The distributions of (D) the chromosome territory strength, defined as the ratio of intra-chromosome to total number of contacts, (E) The radius of gyration, and (F) Inter-chromosome Gauss linking number, are plotted for the different structural variants: SAW, G, Sand R. (G) The quadrants correspond to where the four structural phenotypes arise as we vary short and long-range lengthwise compaction. Within each quadrant, shown are representative snapshots of a chromosome in the left and the genome on the right. The chromosome on the left is colored from blue to white to red from one end to the other. The genome shows five chromosomes in different colors, highlighting the territory strength.

In addition, centromeres and telomeres are incorporated in the model chromosomes, depicting constitutive heterochromatin. The centromere monomers, following the second principle of genome organization, can adhere to other centromere monomers when in proximity. When multiple chromosomes were simulated simultaneously, the self-adhesive interaction was found to drive phase separation of centromeres, leading to spatially segregated clusters of centromeres. Lengthwise compaction of chromosomes was found to establish chromosome territories and screen trans-centromere interactions, counteracting their clustering. Consequently, lengthwise compacted chromosomes showed less-clustered or scattered centromeres. Similar phenomenon is observed for telomere clusters. Note that, by design, this model does not account for features like TADs or A/B compartments, since we are interested in the average features of chromosome organization. Other models have shown that incorporating DNA-sequence-specific heterogeneity, corresponding to active and silent chromatin, in the block co-polymer nature of the chromosomes reproduces TADs and A/B compartments [18, 42, 49].

Next, considering the third principle of structure regulation: interactions of heterochromatin with nuclear lamina, we introduced a confining wall, made up of static monomers, around our multi-chromosomes simulations. The simulated system showed less clustering when the centromeres interacted favorably with the wall.

In our previous work [8], we found that centromeres and/or telomeres of some organisms reside in clusters. Interestingly, the chromosomes of those organisms are likely to not have a fully functional Condensin II, and showed lower territorialization. Moreover, depletion of Condensin II drove a genomic structure with scattered centromeres to the one with clustered centromeres. We used a model similar to the one presented here, where reducing the long-range component of lengthwise compaction led to weaker territories and higher centromere clustering. In this study, we recapitulate the previous result, and analyze in detail the underlying mechanism of screening by chromosome arms in driving the phenomena. We also find that short-range lengthwise compaction is less efficient in screening centromeres and inhibiting centromere clustering, in line with our previous experimental finding that depletion of Condensin I did not affect centromere clustering [8]. Lamina tethering, not analyzed in the previous study, may strongly counteract centromere clustering, independent of lengthwise compaction. Lamina tethering instead of screening centromeres, abolishes contacts with the centromeres by placing them at the nuclear periphery. Indeed, proximity of the centromeres to the wall diminished in experiments showing higher centromere clustering [8].

A specific structural phenotype, showing alignment of chromosome arms on top of each other, we dubbed chromosome “fold over”, was observed in organisms lacking one or more Condensin II subunits [8]. We were unable to structurally identify the fold-over state by just varying lengthwise compaction and centromere phase separation. Here, we find that the respective adhesive interactions of the centromeres and telomeres to the lamina is essential for chromosome fold-over. Moreover, polar lamina interactions, i.e., telomeres and centromeres adhering to opposite hemispheres of the lamina enhances the fold over phenotype. In line with our previous experimental finding, we found that lengthwise compaction, by stiffening the chromosomes, diminished their tendency to fold over.

The model puts forth the idea that a competition among the three generalized forces determines chromosome shape as well as inter-chromosome organization, and provides a conceptual framework to interpret the relationship between activity of various proteins and the genome structure. In future, calibration of this model to specific organisms can identify the relative strengths of the different forces, and may be used to further build on the species-wide structural classification of genomes [8].

## II. METHODS

Our simulations are governed by the stochastic Langevin dynamics:

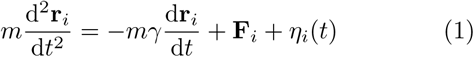

where **r**_*i*_ is the position vector of the *i*-th monomer, *m* = 1 is the mass, *γ* = 0.1 is the friction coefficient, and *η*_*i*_(*t*) is an uncorrelated Gaussian random process such that ⟨*η*_*i*_(*t*)*η*_*j*_(*t*+*s*)⟩ = 2*mγTδ*(*s*)*δ*_*ij*_ (we use reduced units of *k*_*B*_ = 1, and *T* = 120). Finally, **F**_*i*_ is the net thermodynamic force experienced by the *i*-th particle due to its interactions with all other particles in the system, defined as follows.

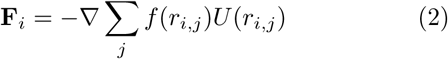

Here, *r*_*i,j*_ is the distance between the particles *i* and *j*, and *f* (*r*_*i,j*_) is the contact function, used to modulate the distance over which inter-particle interactions are effective (see Appendix). This function ensures that the interactions between particles are contact-based and there are no long-range forces in the system [18, 42]. The pairwise interaction potential is given by *U* (*r*_*i,j*_), which only depends on the distance between the two interacting monomers. The interaction potential is written as a sum of different components:

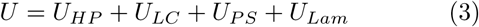

where *U*_*HP*_ is the simple homopolymer potential, representing chromosome beads connected via springs. The other components include lengthwise compaction activity (*U*_*LC*_), phase separation (*U*_*P S*_), and lamina adhesion (*U*_*Lam*_) (see Supplementary materials for details). The lengthwise compaction potential, generating a force that crumples the the polymer along contour length, is implemented as a sum of a short-range and a long-range part. The short-range potential primarily controls the interaction strength between closely spaces monomers along the genomic contour; the long-range controls the interaction strength between loci that are far apart. The phase separation potential stabilizes interactions between certain loci, such as two centromere monomers. The lamina adhesion term stabilizes interactions between the static lamina beads and lamina-adhering beads of the chromosomes. The simulations are performed using the GROMACS package for molecular dynamics [50]. It is important to note that the analyzed simulations trajectories are all in the steady state, and are independent of the initial configurations (Supplementary Materials, see Fig. S13).

## III. RESULTS

### A. Structural diversity driven by differential long- and short-range lengthwise compaction

To understand how lengthwise compaction can modify overall chromosome structure, we used a multi-chromosome simulation setup where we independently varied the intensity of the short- and long-range compaction components of LCP potential and analyzed the structural consequences (Fig. 1).

We found four structural phenotypes of chromosomes depending on the strength of the short- and long-range lengthwise compaction: SAW(self-avoiding walk), Globular (G), String (S), and Rope (R) (Fig. 1E). The SAW chromosomes appeared minimally compact, diffused, and highly entangled or intermingled with each other. The Globular (G) phenotype corresponds to a compact, spherical globule with minimal overlap between adjacent chromosomes. String-like (S) chromosomes appeared as strings that were compact locally but adopted non-compact configurations at the larger scale, and consequently showed some overlap with other chromosomes. Rope-like (R) chromosomes appeared as thick ropes or cylinders, with higher order structures at smaller lengths. R-chromosomes often folded into spheroidal volumes excluding other chromosomes, much like the G phenotype, however, the R phenotype showed a much higher degree of cylindrical organization at smaller lengthscales.

For low overall lengthwise compaction (i.e., both the short- and long-range components are low), LCP model exhibited a self-avoiding random walk (SAW)-like character. SAW chromosomes feature a large radius of gyration, and the probability of contact decays sharply along the genomic contour (Fig. 1A,E). Chromosomes where all SMC complexes are inactive leading to a complete loss of lengthwise compaction are represented by the SAW phenotype.

In presence of lengthwise compaction three other phenotypes emerged. The String phenotype has only short-range compaction, the Globular phenotype has only long-range compaction, while, the Rope-like phenotype has both long and short-range compaction. The probability of intra-chromosome contact for loci pairs that are nearby along the chromosome contour is higher for the phenotypes with strong short-range contacts, i.e. String and Rope-like chromosomes (Fig. 1C). Note, the contact probability between nearest or next-nearest neighbors is controlled by the homopolymer potential (Fig. S2). On the other hand, the contact probability between distant cis-loci pairs is higher for the Globular phenotype. While long-range compaction tends to crumple the chromosomes into a globule, short-range compaction imparts local stiffness such that the chromosomes resist bending that counteracts contacts between distant chromosome loci. This is why the contact probability between distant segments is higher in the Globular state compared to the Rope or in the String compared to the SAW phenotypes (Fig. 1C), possibly providing a mechanical cue via which short-range compaction may antagonize the signatures of long range compaction.

Condensin II is associated with extrusion and stabilization of long chromatin loops [11, 21, 44], and drives the Globular state. While, Condensin I and cohesin, by establishing shorter loops, underlies the Stringy state. Mitotic chromosomes have strong long-range compaction, thanks to the activity of Condensin II, suggesting both the Globular and the Rope phenotypes are possible mitotic structures. Organisms, like yeast [9, 21], containing only one Condensin variant that establishes long-range compaction, exhibit chromosomes with lower length-to-width ratio, consistent with the Globular phenotype (Fig. 1G). While, higher eukaryotes, like humans [9, 44], containing both Condensin variants have both long- and short-range compaction and the mitotic chromosomes have bent rope-like shapes, as seen in the Rope phenotype (Fig. 1G). Interphase chromosomes, on the other hand, have higher short range compaction and modest long range compaction [51], owing to the cohesin and interphase-specific-Condensin II activity. This suggests interphase chromosome shapes are partially globular and territorial due to Condensin II activity, with some locally compact string-like architecture owing to Cohesin activity. Increasing Cohesin residency time on to interphase chromatin has been shown to transform chromosomes into compact mitotic-looking chromosomes in interphase cells, which the authors dubbed the “vermicelli” phenotype [13, 52]. With such a mutation, Cohesin is expected to reinforce both short and long-range compaction and drive Rope phenotypes. Starting from a Rope-like mitotic chromosome, if Condensin I is depleted, we expect a transformation into Globular chromosomes, whereas, if Condensin II is depleted, our model predicts a transition to the Stringy phenotype. Experimental depletion of Condensin I leads to fuzzy-looking, shorter, thicker chromosomes [46, 48], while Condensin II depletion makes chromosomes into thin, cylindrical, string-like objects [46, 48], both inline with our model expectations. The accompanying shift in the probability of contact curves for Condensin I and II depletion [8, 11], are also in general agreement with our model.

### B. Lengthwise compaction establishes chromosome territories and suppresses inter-chromosome entanglements

Chromosome territories are mutually exclusive sub volumes in the nucleus occupied by individual chromosomes [53, 54]. A direct consequence of chromosome territories is a predominance of intra-chromosome contacts over trans-chromosome contacts, hence we define the proportion of intra-chain contacts as a measure of the territory strength. Using Voronoi tessellation to identify contacts between monomers in a structure, we categorize contacts into either intra- or inter-chromosome to calculate the chromosome territory strength (Fig. 1).

Lengthwise compaction is crucial for establishing and maintaining chromosome territories. LCPs show formation of territories of varying strengths depending on the intensity of short- and long-range compaction (Fig. 1D). The long-range compaction component is more effective in establishing contacts between the distant segments of the chromosome, leading to strong territorial chromosomes in the Globular and Rope states, and somewhat weaker territories in the String state. Notably, territories are lost in the SAW state, and chromosomes intermingle with each other (Fig. 1D), leading to weaker intra-chromosome contacts in the contact maps (Fig. S1). This highlights, Condensin II as the major driver of chromosome territories, as has been seen experimentally [8, 54].

Topological constraints, arising from the fact that DNA chains may not spontaneously pass through one another, must be navigated during compaction driven reorganization of the chromosomes. Topology manipulations via chain crossing is mediated by DNA topoisomerase enzymes that are prevalent in the cell [55, 56]. Since these enzymes can only act locally, unaware of the global topology of the chromosomes, we hypothesize that topoisomerases randomly pass strands irrespective of the global chromosome topology [16]. This translates to a topological equilibrium, where inter/intra-chromosome topology (linking number) fluctuates. Our simulations are in a fluctuating topology ensemble, i.e., chains are allowed to cross each other when they collide, albeit overcoming an energy barrier is necessary. The immediate neighbors along the chain experience a mutual hard-core repulsion, whereas, all other monomer pairs (cis and trans) experience a soft-core repulsion upon overlap (Supplementary Materials). Importantly, the soft-core repulsion, though suppresses, does not abolish topology fluctuations. Chain crossing events are essential for dynamics leading to chromosome structure transformations under varied lengthwise compaction.

The inter-chromosome Gauss linking number, representing inter-chromosome entanglements show a broader distribution under lower lengthwise compaction (Fig. 1F). Random collisions between chromosome segments, when leading to a change in topology, may increase or decrease the overall entanglement with equal likelihood. As a result, the distributions are symmetric about zero. The width of the distribution indicates how tightly the inter-chromosome entanglement is regulated. A broad distribution may be inconsequential in some cases, however, a narrow distribution of entanglements is crucial to ensure a faithful disentanglement of mitotic chromosomes. Lengthwise compaction, by reducing inter-chromosome contacts, suppresses inter-chromosome entanglements. Individual topoisomerase enzymes may not sense global entanglement topology of chromosomes, and thus, cannot independently drive chromosome disentanglement. However, topology fluctuations facilitated by the activity of topoisomerase enzymes along with lengthwise compaction by SMC complexes provides an efficient mechanism for driving chromosome disentanglement [16].

### C. Centromere phase separation is counteracted by lengthwise compaction

Self-adhesion between heterochromatin segments leading to their phase separation into three-dimensional compartments is a prominent feature of genome organization [18, 26, 29–32]. So far our LCP model did not have any compartmental segregation forces. To study the interplay of lengthwise compaction with compartmental segregation, we add centromeres into our model. We designated the centrally located polymer block in LCPs as a centromere (constitutive heterochromatin) and added self-adhesive interaction of strength *χ*_*C*_ between the centromere monomers. The magnitude of *χ*_*C*_ represents the energy associated with the interaction between a pair of centromeric loci. We vary *χ*_*C*_ between 0 and -0.3, where *χ*_*C*_ = 0 indicates that the phase-separation based interactions between any two centromeric monomers is the same as that between a non-centromere and a centromere monomer. While, a negative *χ*_*C*_ value indicates that the interaction between two centromere monomers is more favorable. Consequently, *χ*_*C*_ = −0.3 corresponds to a strong favorable interaction between centromere monomers.

Notably, the self-adhesion does not distinguish between cis and trans-centromere monomer pairs. We use a hierarchical clustering algorithm to designate centromeres into spatial clusters, and plot a histogram of the number of clusters corresponding to the ensemble (Fig. 2). We also plot the proportion of trans-centromere contacts, calculated using Voronoi tessellation, that signifies the propensity of inter-centromere interaction (Supplementary Materials). Tendency to phase segregate or compartmentalize should correspond to lower number of clusters and a higher proportion of trans-centromere contacts.

**FIG. 2.**
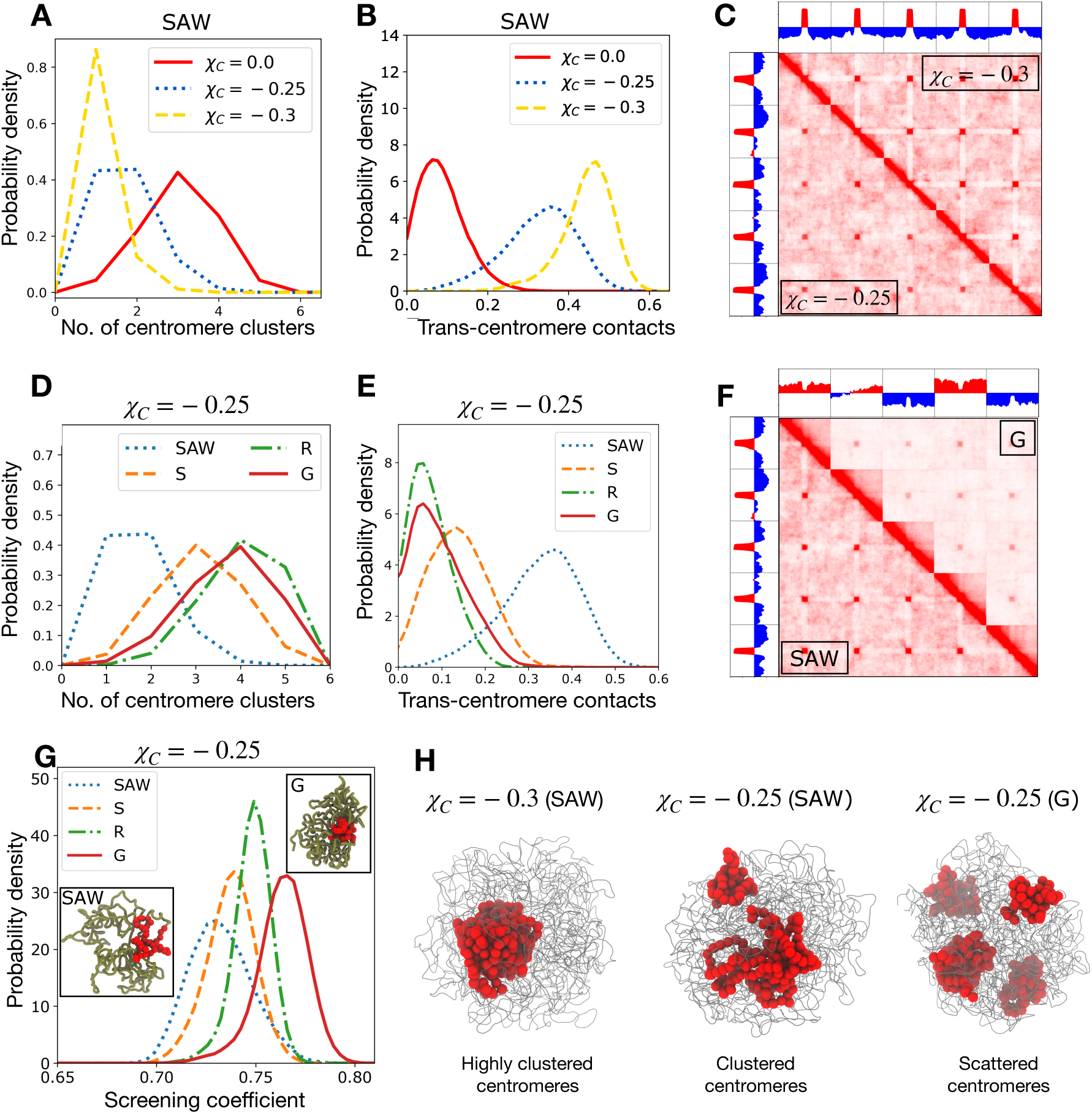
Phase segregation of centromeres and screening of inter-centromere contacts by lengthwise compaction. (A) Probability density of number of centromere clusters, derived from hierarchical clustering, and (B) Proportion of transcentromere contacts, calculated using Voronoi tessellation, plotted for various centromere adhesion strengths *χ*_*C*_ . (C) Contact maps of the genome containing five chromosomes, showing *χ*_*C*_ = −0.3 and *χ*_*C*_ = −0.25 on the upper and lower triangles, respectively. The corresponding principal eigenvectors (PC1) are also shown. Note that centromere clusters form in both the cases, but with a stronger centromere self-adhesion, centromeres are interacting more exclusively with other centromeres. (D) Number of centromere clusters and (E) Proportion of trans-centromere contacts, for a fixed inter-centromere adhesion *χ*_*C*_ = −0.25, and varying lengthwise compaction. (F) Contact map for a fixed centromere self-adhesion (*χ*_*C*_ = −0.25) but different lengthwise compaction (SAW and G-states), showing increase in chromosome territory and lowering of inter-centromere contacts in the G-state compared to SAW. The corresponding PC1s are shown. (G) The screening coefficient, defined as the ratio of contacts between a centromere and its chromosome arms to all the contacts made by the centromere. The screening is higher for higher lengthwise compaction (especially, the long range component), a consequence of burying centromeres within the chromosome territory. (H) Simulation snapshots showing the entire genome as thin gray tubes and centromeres as red spheres.

When the centromere adhesion is strong (*χ*_*C*_ = −0.3), there are abundant trans-centromere contacts, leading to a global phase-separation and the centromeres form one macro cluster (Fig. 2A,B). Lowering the magnitude of *χ*_*C*_ leads to a higher average number of clusters, lower proportion of trans-centromere contacts, and a lower tendency to phase segregate. The case with no centromere attraction (*χ*_*C*_ = 0) shows the least tendency to cluster, and the centromeres are peripherally located. The origin of this behavior is likely entropic, as has been observed in previous models [58]. The macro-phase separated centromere clusters are more compact and reside in the interior of the nucleus for strong adhesive interactions (Fig. 2).

Centromeric clusters appear as “dots” or focal interactions in the inter-chromosomal region of the simulated contact maps (Fig. 2C). Higher self-adhesion increases the specificity of centromere contacts, i.e., the centromere predominantly interacts with other centromeres instead of the rest of the chromosomes. This results in the appearance of white stripes in the intra-chromosomal contact maps for *χ*_*C*_ = −0.3 (Fig. 2C).

Higher clustering for increased self-adhesion of centromeres occurs irrespective of lengthwise compaction (Figs. S3-S5). Interestingly, for a fixed intensity of centromere adhesion, we find that the number of centromere clusters are the lowest in the SAW state. Whereas, the clustering tendency of centromeres is lower, i.e., the number of centromere clusters are larger, when the chromosomes are lengthwise compacted (Fig. 2D). This suggests a role of lengthwise compaction in inhibiting centromere clustering. Lengthwise compaction establishes chromosome territories that bury the centromeres and screen the inter-centromere (trans-chromosomal) interactions, thus counteracting their tendency to phase segregate. We define a screening coefficient as the ratio of the number of contacts a centromere monomer makes with the flanking arms of the chromosome, to the total number of contacts of the monomer. Consequently, when the centromere interacts strongly with the cis-chromosome arms, the screening coefficient is higher and inter-centromere contacts are depleted. Figure 2G shows the screening coefficient is higher for structures with higher lengthwise compaction, especially the long-range component. Also note, the drastic decrease in contacts in the inter-chromosomal region of the contact map for the Globular state, along with a lower probability of inter-centromeric contacts (Fig. 2), which is a direct consequence of the screening of inter-chromosome contacts by lengthwise compaction. The antagonistic behavior between Condensin II activity and centromere clustering has been confirmed experimentally [8].

The principal eigenvectors (PC1) of the correlation matrix associated with intra-chromosomal contact maps is typically associated with chromosome compartments [2, 18]. PC1 calculated from our full genome contact map encodes information related to both chromosome territories and centromere clusters (Fig. 2C, F). PC1 for the less compact (SAW) state with centromere clusters, has opposite signs corresponding to chromosome arms and centromeres, reflecting a boundary between the clustered centromere phase and the rest of the genome. On the other hand, PC1 for the Globular (G) state, instead of changing signs at centromeres, mainly changes sign corresponding to change from one chromosome to another, i.e., at the territory boundaries. Centromere clusters dominate in the less-compact state, whereas, chromosome territories dominate in the states with higher lengthwise compaction states, which reflects in the structure of the corresponding PC1.

Adhesive interaction between telomeres leads to similar phenomena where telomeres tend to form clusters that are counteracted by lengthwise compaction via screening inter-telomere interactions (Fig S6-S7). However, telomeres being larger in number than centromeres have higher entropy and show a lower tendency to phase segregate compared to centromeres.

### D. Lamina tethering of centromeres counteracts their clustering

Physical tethering of heterochromatin domains to the nuclear envelope or lamina, mediated by the lamin proteins, is a well known aspect of nuclear organization of the genome [29, 33–37, 39]. To model the lamina, we placed static beads covering the nuclear periphery and introduced adhesion between centromeres and a subset of static lamina beads (Supplementary Materials). The adhesive interaction was implemented using the same procedure as for centromere adhesion. We introduced *χ*_*L*_, which parameterizes the interaction between centromeres and the sticky lamina beads, where *χ*_*L*_ = 0 indicates no adhesion between the centromere and the lamina (Fig. 3). While, *χ*_*L*_ < 0 indicates a favorable interaction between the centromere and the lamina, and a higher negative value represents a stronger adhesive strength. The adhesive interaction was found to drive the centromeres to the nuclear periphery (Fig. 3C, S10). This geometric reorganization of the genomic structure upon lamina interaction has been observed in other modeling approaches [29, 39], however, what role it plays for centromere clustering is less clear.

**FIG. 3.**
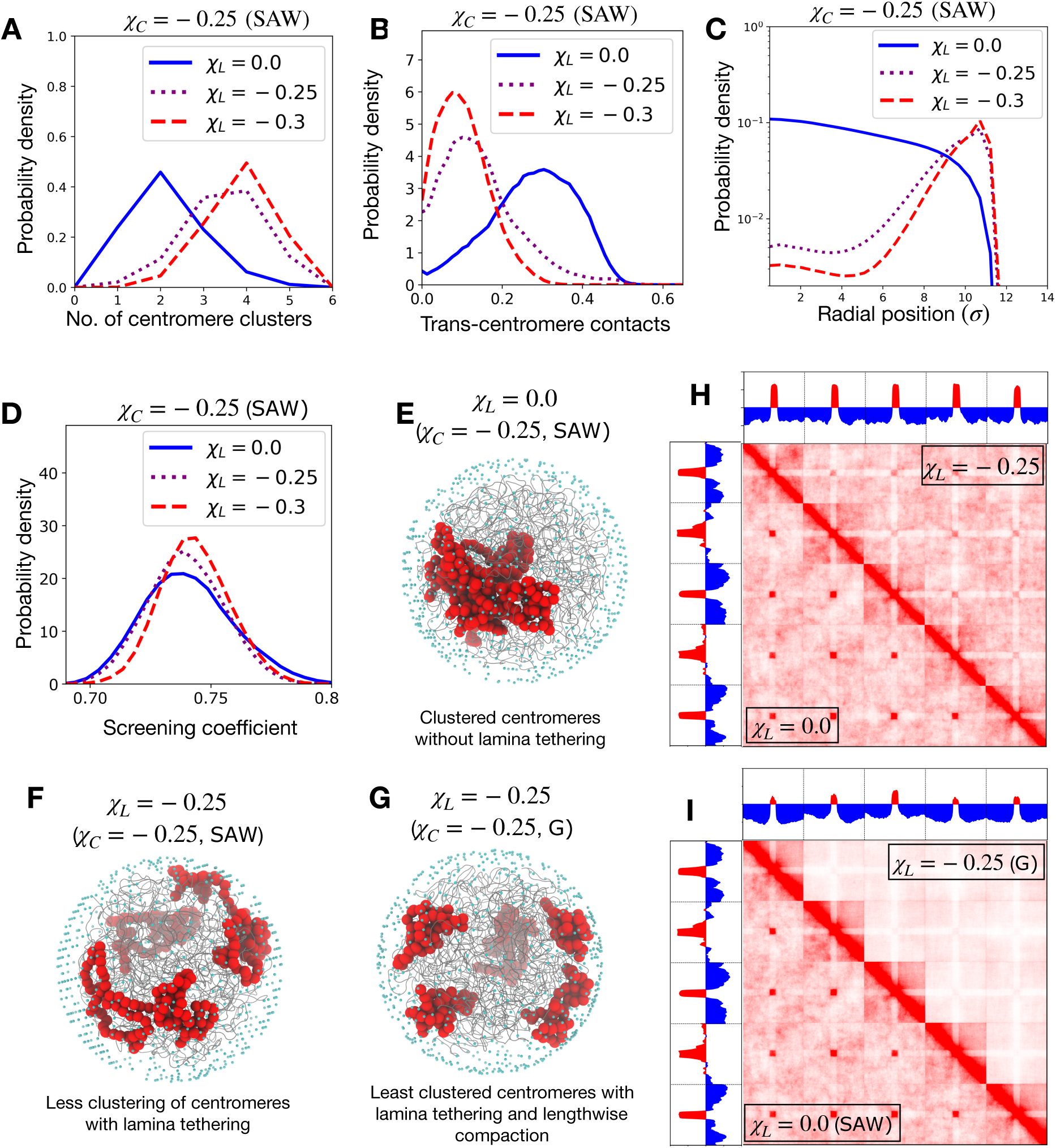
Lamina tethering counteracts segregation of centromeres. (A) Probability density of the number of centromere clusters, (B) proportion of trans-centromere contacts, and (C) radial distribution of centromeres, plotted for various strengths of lamina-centromere adhesion *χ*_*L*_. The centromere self-adhesion is fixed *χ*_*C*_ = − 0.25, as well as the lengthwise compaction state (SAW). (D) The distribution of screening coefficients for different lamina adhesion strengths, with a fixed centromere self-adhesion (*χ*_*C*_ = − 0.25) and fixed lengthwise compaction (SAW). (E), (F), and (G) Simulation snapshots showing the chromosomes as gray tubes, the centromeres as big red spheres, and the lamina beads as small cyan spheres. (H) Contact maps for genomes containing five chromosomes with the same centromere self-adhesion (*χ*_*C*_ = − 0.25) and same lengthwise compaction (SAW), but different lamina adhesion of centromeres: the upper triangle is for the case of centromeres adhering to lamina (*χ*_*L*_ = − 0.25), while lower triangle is without lamina adhesion (*χ*_*L*_ = 0.0). (I) Genome contact map for five SAW chromosomes with centromere adhesion without lamina tethering(*χ*_*C*_ = − 0.25, *χ*_*L*_ = 0.0) is shown in the lower triangle, while the upper triangle shows lengthwise compacted Gchromosomes with centromere adhesion as well as lamina adhesion of centromeres (*χ*_*C*_ = −0.25, *χ*_*L*_ = −0.25).

Lamina tethering of centromeres led to a decreased clustering tendency of centromeres. For a fixed inter-centromere adhesion, increasing lamina-centromere adhesion (*χ*_*L*_ = − 0.25 or − 0.3) decreased trans-centromere contacts, increased the number of clusters, and the corresponding contact maps showed lower interactions between centromeres of different chromosomes (Figs. 3A,B,F).

The role of lengthwise compaction in counteracting clustering of centromeres remains unaltered in presence of lamina tethering (Fig. S9-S10). Since both lamina interaction and lengthwise compaction independently counteract clustering of centromeres, formation of globally segregated centromeres is strongly attenuated when the chromosomes are lengthwise compacted in presence of lamina-centromere adhesion (Figs. 3H, I and Fig. S8). However, the mechanisms underlying inhibition of centromere clustering by lamina adhesion and by length-wise compaction are different: the screening coefficient does not increase with higher lamina interactions (compare Figs. 3D and 2G). Lengthwise compaction enhances intra-chromosome interactions between the centromere and the chromosome arms that leads to a reduction of inter-centromere contacts. Whereas, lamina isolates the centromeres to the periphery, thus generally inhibiting all contacts with the centromeres. The depletion of contacts between centromere and chromosome arms for strong lamina tethering underlies the white stripes in the intra-chromosomal maps, and a different structure of the PC1 (compare Figs. 3I and 2F). Many genes or gene clusters are preemptively released from the lamina during cell differentiation that are upregulated in the following stages [34]. This isolation of genomic elements due to lamina tethering likely contributes to the maintenance of a transcriptionally silent state. While the release from lamina favors contacts between the released segment and its potential regulatory elements, thus making the gene accessible to the transcriptional-regulation machinery.

In our previous work [8], wild-type human genome showed scattered centromeres that resided near the nuclear lamina and had strong lengthwise compaction (and consequently, well defined territories). While, the Condensin II-depleted phenotype showed clustered centromeres that moved to the interior of the nucleus. Our results suggest that both lengthwise compaction by Condensin II and lamina tethering are important forces responsible for scattered centromeres in wild-type cells. The clustered centromere phenotype is characterized by the loss of both these clustering-inhibitory forces. The model further predicts a less strong phenotype, i.e., less clustering of centromeres, if only one the forces, either lamina tethering or lengthwise compaction is depleted keeping the other same.

### E. Chromosome “fold over” is facilitated by strong clustering of centromeres and telomeres along with their tethering to the lamina

Chromosomes in some organisms, like bread wheat and yellow-fever mosquito, are known to adopt a configuration where the chromosome arms align on top of each other so as to form a telomere-to-centromere axis, which we previously called the “fold-over” phenotype [8]. These organisms are likely to not have all the Condensin II subunits, i.e., they likely lack long-range lengthwise compaction [8]. In line with this, we find that higher lengthwise compaction enhances the chromosome bending stiffness, which straightens the chromosome axis and counteracts fold over.

To quantify chromosome fold-over in structural ensembles, we considered the centroid of each 100 consecutive monomers, giving an average shape of the chromosomes (Fig. 4A), and then computed the bend angles at the centromere (*θ*_*centro*_) and the arms (*θ*_*arms*_). Fold-over configuration requires a low *θ*_*centro*_ and a high value of *θ*_*arms*_. Lengthwise compaction tends to the straighten the chromosomes resulting in a lower value for both the angles (Figs. 4B,C). Adhesion of centromeres or telomeres leading to their clustering has a weak effect on fold over. However, lamina adhesion of the centromeres and telomeres, along with their strong respective clustering tendency did reproduce the fold-over phenotype (Fig. 4D). Strong lamina tethering of centromeres and telomeres compete with lengthwise compaction, weakening the shifts in *θ*_*centro*_ and *θ*_*arms*_ with lengthwise compaction. When centromeres and telomeres interact strongly with the lamina, it is the cylindrical, length-wise compacted state (e.g., Rope-like phenotype) that produced the stronger fold-over signal (Fig. S11-S12).

**FIG. 4.**
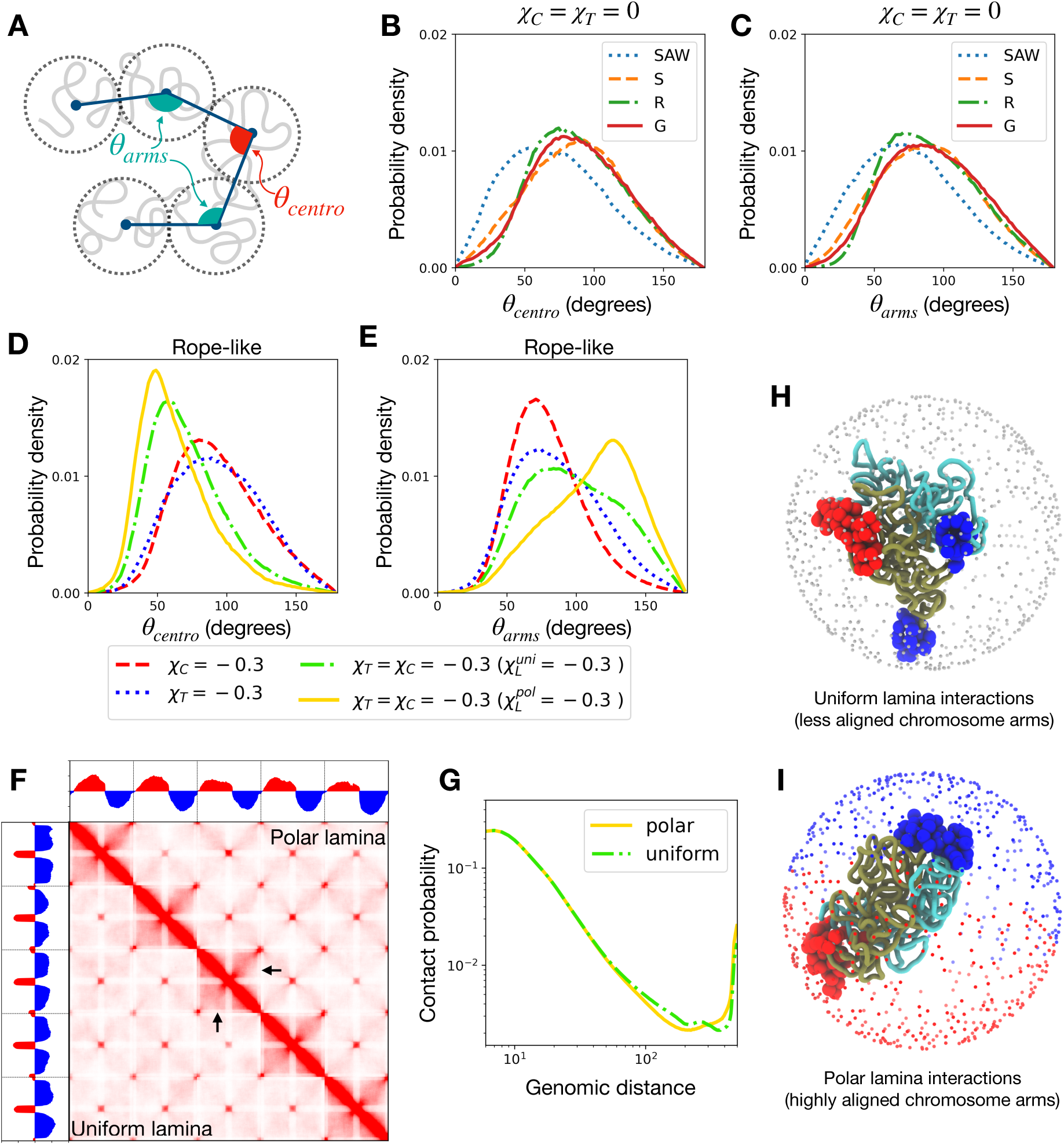
Chromosome fold over is facilitated by strong centromere and telomere clustering and lamina adhesion. (A) Schematic of chromosomes showing the bend angles characterizing the fold over: *θ*_*centro*_ and *θ*_*arm*_. Folding over requires a low *θ*_*centro*_ and high *θ*_*arm*_. (B) and (C) Probability distributions of the characteristic bend angles are plotted for various lengthwise compaction without any centromere or telomere adhesion. The bending of the chromosomes at the centromere as well as at the arms are in general counteracted by lengthwise compaction. (D) and (E) Probability density of the bend angles in the Rope-like (R) state, under various conditions such as centromere (or telomere) phase separation, and adhesion with uniform or polar lamina. Polar lamina produces the strongest fold over signal. (F) Contact maps of the genome with five Rope-like chromosomes with strong centromere and telomere adhesion (*χ*_*C*_ = *χ*_*T*_ = − 0.3) along with their uniform adhesion to the lamina 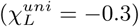 shown in the lower triangles, while the upper traingle corresponds to a polar lamina configuration 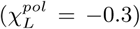. The black arrows show the region where there is loss of telomere to centromere contacts in the polar case. (G) Chromosome contact probability curves corresponding to the polar and uniform cases in the contact map. (H) and (I) Simulation snapshots showing one chromosome with centromeres as big red spheres and telomeres as big blue spheres, and the two chromosome arms are shown as dark green and cyan tubes. For the uniform lamina case (H), the lamina beads are shown as small gray spheres, while for the polar case, the lamina beads interacting favorably with the centromere are shown as small red spheres and that with the telomere are shown as small blue spheres.

Introducing a polar geometry in the lamina adhesion mechanism, i.e., centromeres and telomeres favorably adhere to distinct hemispheres, was found to reinforce the fold-over phenotype (Figs.4D,F). In addition to lowering the angle at the centromere, polar lamina interaction straightened the arms of the chromosomes, enhancing the centromere-to-telomere axis (Figs. 4E, I). The straightening of the chromosome in polar lamina led to lower contact probability between distant cis chromosome segments (Fig. 4H).

The signature of the fold-over in the contact probability matrices is that of a characteristic counter-diagonal in the intra-chromosome blocks (Fig. 4G) [8]. The inter-chromosome blocks of the contact matrices show characteristic X-shaped patterns corresponding to the alignment of inter-chromosome arms, and intense focal interactions corresponding to the clustering of centromeres and telomeres. PC1 of the two contact matrices corresponding to uniform and polar lamina show very different structures. The PC1 for uniform lamina interactions changes sign corresponding to centromeres and telomeres, indicating formation of the respective compartments (Fig. 4G). For polar lamina interactions, PC1 shows a unique undulatory structure where the two arms have different signs.

The model overall suggests chromosome fold over is expected to be most prominent in genomes featuring strong telomere and centromere clustering and adhesion of the respective clusters to a polar lamina configuration. Recent data-driven modeling of the yellow-fever mosquito genome, which shows fold-over chromosomes, are in agreement with these findings [42].

## IV. DISCUSSION

We simulated a genome consisting of multiple coarsegrained chromosomes, where we investigated the role of three fundamental forces involved in genome organization: lengthwise compaction, phase separation, and lamina tethering of chromosomes. We started with modeling chromosomes as a lengthwise-compacted polymer (LCP), that incorporates the steady-state activity of SMC complexes via a LCP potential capable of compacting the chromosomes in a lengthwise manner. Inspired by the differential activity of SMC variants (Condensins and Cohesin), we decomposed the LCP potential into two parts: one that locally compacts the chromosome polymer (short-range), and the other brings together distant parts of the chromosome (long-range). We found that the short- and long-range components of length-wise compaction have distinct phenotypical consequences on the chromosome structure (Fig. 1). While reduction in lengthwise compaction led to a loss of chromosome territories and intermingling of chromosomes, the long-range potential drove territorial, globular chromosomes, and the short-range compaction component led to thin, cylindrical, string-like chromosomes. Chromosomes with both long- and short-range compaction, as occurs in mitosis of higher eukaryotes, exhibit thick rope-like shapes. The transformations from one structural phenotype to another is consistent with experimental observations of chromosome shapes in conditions of SMC depletion [13, 46, 48, 52].

Importantly, the results rationalize our previous experimental finding that organisms lacking SMC subunits corresponding to Condensin II, or cells following depletion of Condensin II [8, 54], have weaker chromosome territories. Condensin II has been implicated in a variety of seemingly unrelated phenomena, such as, inhibition of chromosome translocation [47], pairing of homologous chromosomes [59], regulation of cellular senescence [60, 61]. We think that Condensin II-driven depletion of trans-chromosome interactions via long-range lengthwise compaction, contributes to these phenotypic occurrences, since all these processes strongly depend on the propensity to form trans-chromosomal contacts.

An important prediction of our model is lengthwise compaction driven territories are maintained even in the presence of topoisomerase-mediated chain crossing. This complements findings that a bead-spring description of chromosomes (without lengthwise compaction) shows a gradual loss of territories [62]. Our model also suggests that the lengthwise compaction activity of SMC complexes may play a direct role in the regulation of inter-chromosome entanglements (Fig. 1). Lengthwise compaction, by reducing inter-chromosome contacts, inhibits the topoisomerase-introduced topology fluctuations from increasing inter-chromosome entanglements [16]. This is supported by *in vitro* experiments reporting the necessary presence of both DNA topoisomerase and SMC complexes in order to individualize chromatids [63]. Synergistic coordination between topoisomerases and SMC complexes at the molecular level has also been proposed, but remains to be validated experimentally [64].

Following the second principle of genome organization, inter-centromere adhesion, corresponding to self-adhesion among heterochromatin, drove the centromeres to phase segregate into large clusters (Fig. 2). Screening of trans-chromosomal contacts by lengthwise compaction impedes global phase segregation of centromeres, and instead there is micro-phase separation into multiple smaller clusters (Fig. 2). In agreement with this, we recently reported experiments where depletion of Condensin II transformed a wild-type nucleus with microphase separated centromeres into a global, macro-phase separated phenotype [8]. Further, a recent study reported increase in compaction and decrease in inter-centromere contacts upon depleting HP1, a protein responsible for phase segregating heterochromatin [28]. As the authors found via modeling [28], only including phase-separation-driving terms is not enough to recapitulate the increase in compaction. Within our framework of three fundamental forces, depletion of HP1 reduces the intensity of self-adhesion, which makes lengthwise compaction by SMC proteins the dominant force, rationalizing the increased intra-chromosome compaction. The competition between lengthwise compaction and phase separation has been hypothesized in other models of genome organization, where simulations of loop extrusion was observed to counteract compartmental segregation, mainly derived from the nonequilibrium nature of loop extruding factors [65]. Our model argues there is also an effective equilibrium description where there is steady-state screening of genomic interactions, especially the inter-chromosome ones, by chromosome territories established via lengthwise compaction.

We have simplified the sequence complexity of the genome in this manuscript, as such, our model lacks the typical sequence heterogeneity that gives rise to transcriptionally active or silent regions along the chromosomes. Heteropolymer models implementing different blocks derived from histone marks corresponding to regions such as euchromatin, heterochromatin, and nucleolus-interacting regions [18, 66, 67], have suggested that nuclear compartments [2] are a result of phase separation of blocks. We note that the underlying principle of phase separation in block copolymer models is the same that drives phase separation of centromeres in our model. Hence, the antagonistic behavior observed between centromere clustering, and lengthwise compaction or lamina tethering may be extended to compartments. In accord with is expectation, antagonism between SMC activity and compartmentalization has been observed experimentally [22, 65].

When nuclear lamina was introduced the confinement did not alter the structures, since the generic adhesive interactions between all monomers were enough to maintain the physiological volume fractions in our setup. However, introducing adhesion of centromeres to the lamina led, not only to geometric rearrangement of centromeres to the periphery but also counteracted their macro-phase separation (Fig. 3). Lamina tethering brings the centromeres to the surface of the nucleus, and in doing so, dissociates the macro-cluster. In absence of favorable lamina interactions, the macro-phase segregated cluster prefers to reside inside the nucleus (Fig. 2). This is consistent with repressive chromatin compartments lying within the chromosome territories [68], and the “inverted” geometry of the heterochromatin in lamin-depleted nuclei [29, 36]. Based on our model results, we expect the “inverted” nuclei phenotype to have a lower lengthwise compaction or a steeper contact probability decay, as was observed [29]. The model further predicts that sufficiently increasing lengthwise compaction in the “inverted” nuclei will perturb the macro-phase segregated heterochromatin cluster, leading to a microphase separated state.

Centromere clustering was strongly inhibited when lamina tethering and lengthwise compaction were both present (Fig. 3). In experiments with Condensin II depletion [8], the centromeres were found to move away from the lamina, which aided their macro-phase segregation. Interestingly, the lamina-release of centromeres was triggered by Condensin II depletion and without any targeted depletion of lamina-tethering proteins [8].

Chromosomes in some organisms have been shown to assume a folded-over structure, where the chromosome arms align such as to form a centromere-to-telomere axis [8]. The model suggests, strong respective clustering among centromeres and telomeres, along with their tethering to lamina is essential for the folded over chromosomes (Fig. 4). When centromeres and telomeres favorably interacted with the opposite poles of the lamina, the fold-over phenotype was enhanced. Interestingly, when clustering of centromeres and telomeres is weak, length-wise compaction counteracted fold-over, however, under strong self-adhesion of centromeres and telomeres, the lengthwise compacted state generated better resolution of the centromere-to-telomere axis. In line with this, recent modeling of the Mosquito genome, which shows the fold-over phenotype, shows that a close agreement with the experimental HiC maps is possible when centromeres and telomeres are confined to the opposite poles of the nucleus and stretched, mimicking lamina tethering [42].

In all, we simulated a model genome and studied the essential physical forces underlying genome organization: lengthwise compaction, self-adhesion of chromosome monomers (centromeres, and telomeres), and tethering of chromatin to lamina. These forces effectively capture the structure of the genome sculpted by a variety of proteins, including SMC complexes and chromatin remodeling proteins that modify the epigenetic landscape of chromosomes. We show that the interplay of these forces can lead to qualitatively different structures, with consequences leading to distinct experimental signatures.

## V. ACKNOWLEDGEMENTS

This work was supported by the Center for Theoretical Biological Physics sponsored by the National Science Foundation (Grant PHY-2019745), by the Welch Foundation (Grant C-1792), and by NSF-CHE-1614101. JNO is a CPRIT Scholar in Cancer Research.

## Supplementary Material

### I. SIMULATION METHODS

#### Contact function

The contact function is a sigmoidal function that cuts off interactions between distant monomers, and this is used to calibrate the contact probability between genomic segments [1]. The contact function *f* (*r*_*i,j*_) is defined as:

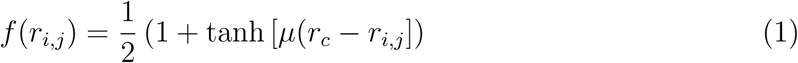

where *µ* = 3.22 ad *r*_*c*_ = 1.78 are used following previous calibration with experimental Hi-C maps [1]. Note, the qualitative results discussed in the main text are not sensitive to small changes in these parameters.

#### Interaction potential

##### Homopolymer

The homopolymer potential (*U*_*HP*_) models a generic bead-spring polymer in which each bead represents a genomic segment containing 20-50 Kb of DNA, where chromosome topology fluctuations are controlled by using an energy barrier. This potential consists of the following five terms, *U*_FENE_, *U*_Angle_, *U*_*hc*_ and, *U*_*sc*_:

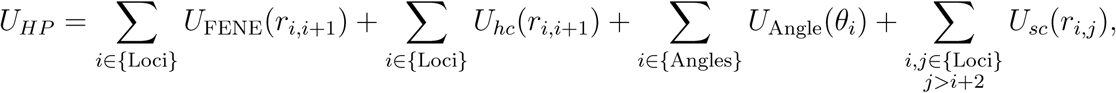

where,

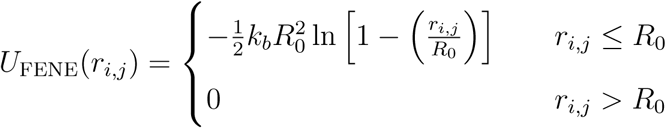

*U*_FENE_ (Finite Extensible Nonlinear Elastic potential) is the bonding term applied between two consecutive monomers of a chromosome.

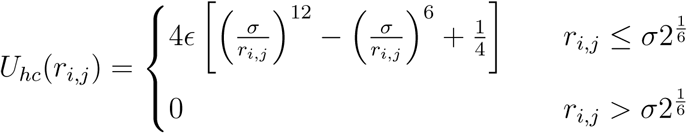

*U*_*hc*_(*r*_*i,j*_) is the hard-core repulsive potential, include to avoid overlap between the bonded nearest neighbor monomers.

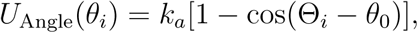

a three-body potential applied to all connected three consecutive monomers of a chromosome, where Θ_*i*_ is the angle defined by two vectors 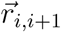 and 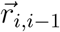, and *θ*_0_ = 0 is the equilibrium angle.

The non-bonded pairs is defined by a soft-core repulsive interaction in the following form:

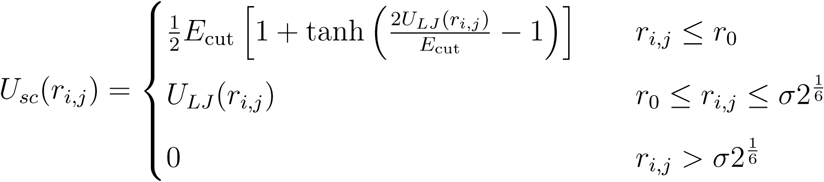

The expression *U*_*LJ*_ correspond to the Lennard-Jones potential:

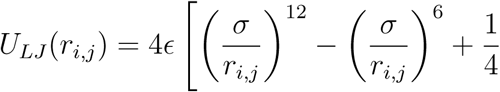

capped off at a finite distance, thus allowing for chain crossing at finite energetic cost. The parameter *r*_0_ is chosen as the distance at which 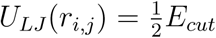. Note that this potential is applied across all non-neighboring monomers of the system.

##### Lengthwise Compaction

We implement lengthwise compaction of the polymer as a sum of two terms:

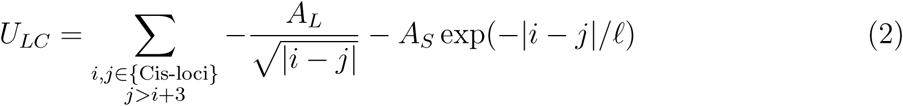

where *ℓ* = 10 is the characteristic length of short-range compaction. The potential is applied between intra-chain monomers that are more than 3 monomers apart, and underlies a looping tendency. The first term with amplitude *A*_*L*_ > 0 controls the long range compaction, while the second term with amplitude *A*_*S*_ > 0 controls the short-range compaction. Note that the intensity of lengthwise compaction depends on the genomic distance between the two loci, and that this potential does not act across chromosomes. Different values of *A*_*L*_ and *A*_*S*_ leads to the structural phenotypes described in the main text: SAW (*A*_*L*_ = 0.05, *A*_*S*_ = 0.05), Globular (*A*_*L*_ = 0.4, *A*_*S*_ = 0.05), String-like (*A*_*L*_ = 0.05, *A*_*S*_ = 0.4), and Rope-like (*A*_*L*_ = 0.4, *A*_*S*_ = 0.4).

##### Phase Separation

The potential associated with phase separation is self-adhesion among monomers. There is a generic adhesion between any two monomers of intensity *χ* = −0.2. This implies whenever two monomers come within interaction distance of one another the energy of the system lowers by *χ* (where the units are in simulation energy scale *ϵ* = *k*_*B*_*T*). The centromere monomers adhere to other centromere monomers with an enhanced adhesive interaction *χ*_*C*_. The interaction potential is represented as follows:

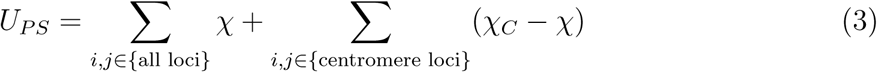

##### Lamina Adhesion

We place static monomers at a distance *R*_0_ from the center of mass of the genome, such as to form a rigid spherical shell of radius *R*_0_ encapsulating the genome. The numerical value of *R*_0_ is decided from the requirement of physiological volume fraction (*ϕ* ≈ 0.1) of the genome inside the nucleus: *R*_0_ = *σ*(*N/*(8*ϕ*))^1*/*3^, where *N* is the total number of monomers in the genome, and *σ* = 1 is the monomer diameter.

While all the lamina beads and genome beads experience a soft-core repulsion, a randomly selected subgroup of 30% of the surface beads interact favorably with the centromere with an interaction strength *χ*_*L*_ *≤* 0. The nuclear envelope contains many elements like the nuclear pores that cover a significant portion of the surface area that are not adhesive to the chromosome segments. Given the genome volume fraction is about 10%, using a sub population of surface beads made the competition between phase separation and lamina adhesion occur for similar values of *χ*_*L*_ and *χ*_*C*_. The lamina interaction potential may be expressed as follows:

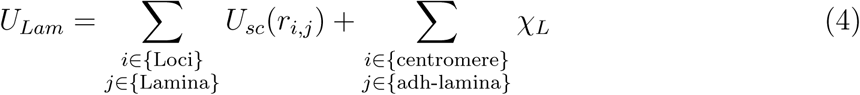

where ‘adh-lamina’ refers to the subgroup of lamina monomers that adhere to the centromere.

### II. TRAJECTORY ANALYSIS

The analysis were done on an ensemble of simulated trajectories. We simulated each parameter set for 2 × 10^7^ time-steps (*dt* = 10^−3^), and generated 10 replicas of each trajectory with randomized initial configurations. We use high temperature annealing of the homopolymer model to generate many random structures that are used to initialize the simulations. We then neglect the initial 10^6^ time steps from our analysis, to ensure the steady-state nature of our trajectories (Fig. S13).

#### Contact probability matrices

Contact probability matrices, the analogue of HiC-maps, are calculated using the contact function. For every frame, we compute the pairwise distance between monomers and then use the contact function to convert the distance into contact probability, following our previous approach [1]. All the snapshots corresponding to a parameter set are then averaged to generate the contact maps shown in the main and supplementary figures.

##### Principal component eigenvector

The outer product of the contact probability matrices were used to generate the correlation matrix. The eigenvector corresponding to the largest eigenvalue is the principal component eigenvector shown along with the contact maps. These principal components have been used to annotate compartments in HiC experiments [1, 2].

#### Voronoi tessellation: Territory strength and trans-centromere contacts

One snapshot of the trajectory can be considered to be a discrete distribution of points in 3D space. Using Voronoi tessellation, the empty space between points maybe filled by non-overlapping polyhedrons, each encapsulating one bead. Each surface of a polyhedron is such that it is a plane perpendicularly bisecting the line connecting a bead to its neighbor. The number of surfaces of the polyhedron defines the number of neighbors for the encapsulated bead. We use the python package: Scipy.spatial.Voronoi [3] to compute the Voronoi diagram in our simulated trajectories.

Using this scheme, we identify neighbors of a monomer, and then classify the ratio of number of intra-chain contacts to the total number of contacts per chain as the strength of the chromosome territory. Similarly, the proportion of trans-centromere contacts is computed from the ratio of number of inter-chromosome centromere contacts to the total number of centromere-centromere contacts.

#### Shape analysis: Gyration tensor

Attributes of chromosome shape, like the radius of gyration and the shape anisotropy, are calculated using the gyration tensor *G*, defined as follows for a set of coordinates:

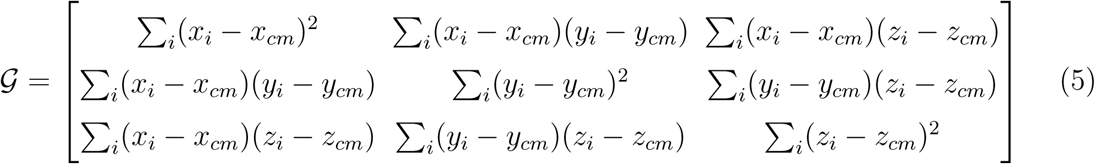

Here the *i*-sum is over all the monomers of the polymer whose shape we are interested in, and (*x*_*cm*_, *y*_*cm*_, *z*_*cm*_) is the center of mass of the polymer. Once this matrix is computed for a given snapshot, we compute the eigenvalues of the gyration tensor *λ*_1_, *λ*_2_, *λ*_3_ (note, *λ*_1_ *≥ λ*_2_ *≥ λ*_3_ > 0) and then calculate the shape descriptors as follows [4]:

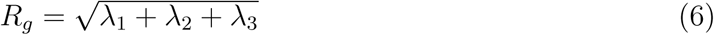

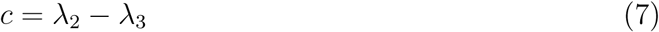

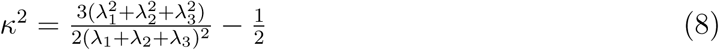

where *R*_*g*_ is the radius of gyration. *c* is the acylindricity, which is lower if there is cylindrical symmetry in the conformation. And, *κ*^2^ is the relative shape anisotropy that is bound between 0 and 1, and is higher for anisotropic structures.

#### Hierarchical clustering: Number of centromere clusters

Clusters of centromeres and telomeres were defined using the hierarchical clustering algorithm via constructing a dendrogram. At the first step, each centromere monomer is considered a cluster, this is the largest possible number of clusters in the genome. Iteratively, clusters are merged, following a condition that the shift in the centroid of the cluster due to the merge is smaller than a cut-off value. We choose this cut-off to be twice the radius of gyration of the cluster (note, using a slightly different value, like three times the radius of gyration, does not change the qualitative nature of our results). When merging two clusters is shifts the centroid to larger than the cut-off, those two clusters are identified as two individual clusters. We use the python module Scipy.cluster [3] to implement hierarchical clustering. The number of clusters obtained from every snapshot is then plotted using a histogram.

#### Radial density distribution

Radial distribution of monomers is calculated from the snapshots. The volume occupied by the genome is divided into concentric shells centered at the centroid of the genome, then the number of monomers is each cell is counted. To obtain the density, the number is divided by the volume of the shell.

#### Gauss linking number: inter chromosome entanglements

The Gauss linking number between two chromosomes, counts the number of signed crossings between the them, and measures their entanglement. We use the “method 1a”, as prescribed in Ref. [5], to compute the linking number. Since linking number is defined only for closed curves, we simply connect the two ends of each chromosome to close the curve during our computation. We calculate entanglement for each snapshot of an ensemble and then plot the histogram of linking number values, showing the distribution of linking numbers.

#### Simulation snapshots

The simulation snapshot images are made using the VMD software [6].

## III. SUPPLEMENTARY FIGURES

See Figures S1-S13.

**FIG. S1.**
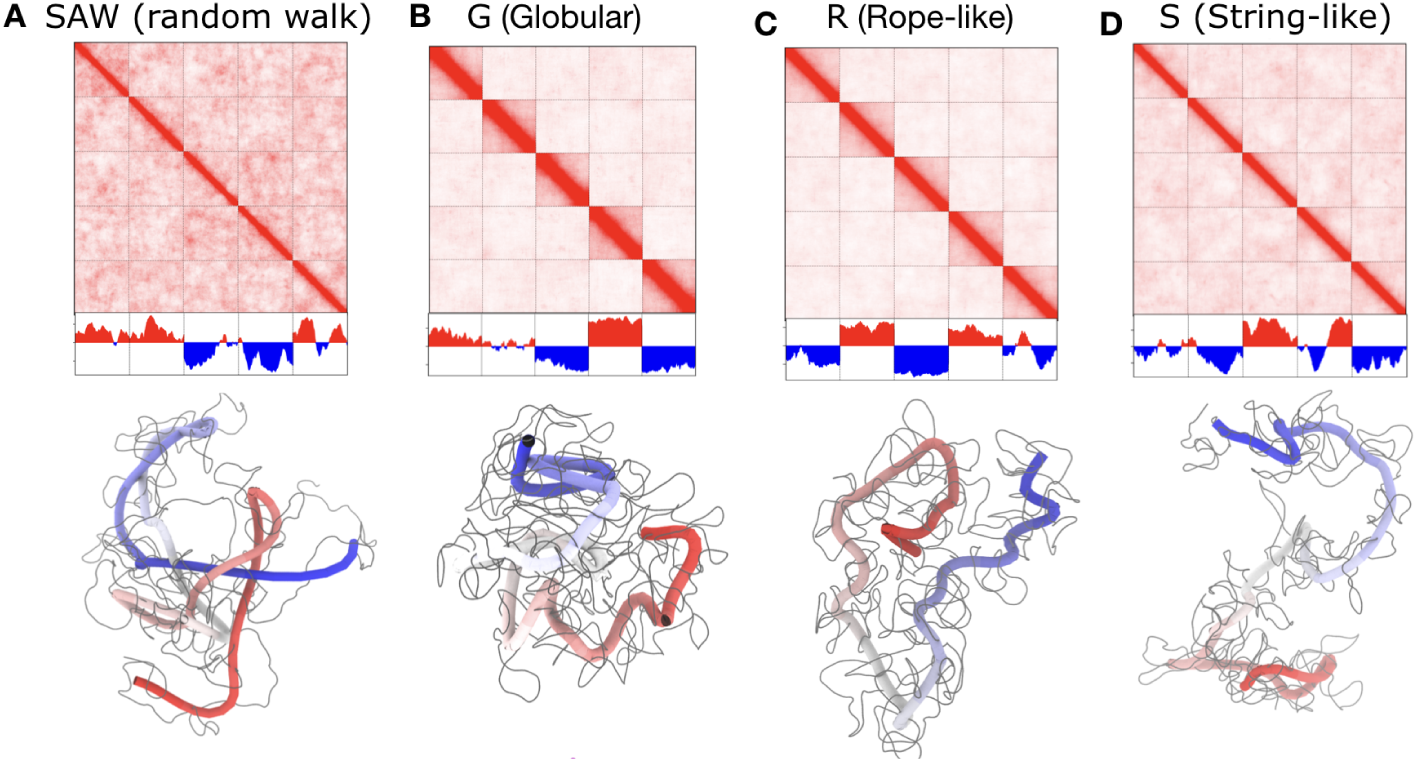
Lengthwise compaction controls chromosome territories. Contact maps corresponding to (A) SAW, (B) Globular (C) Rope, (D) String phenotypes are shown for chromosomes with *χ*_*C*_ = *χ*_*T*_ = 0. Below each contact map are representative structures. A chromosome when renormalized by 20 monomers, shows the underlying backbone. This backbone is shown as a tube in blue-to-red coloration.

**FIG. S2.**
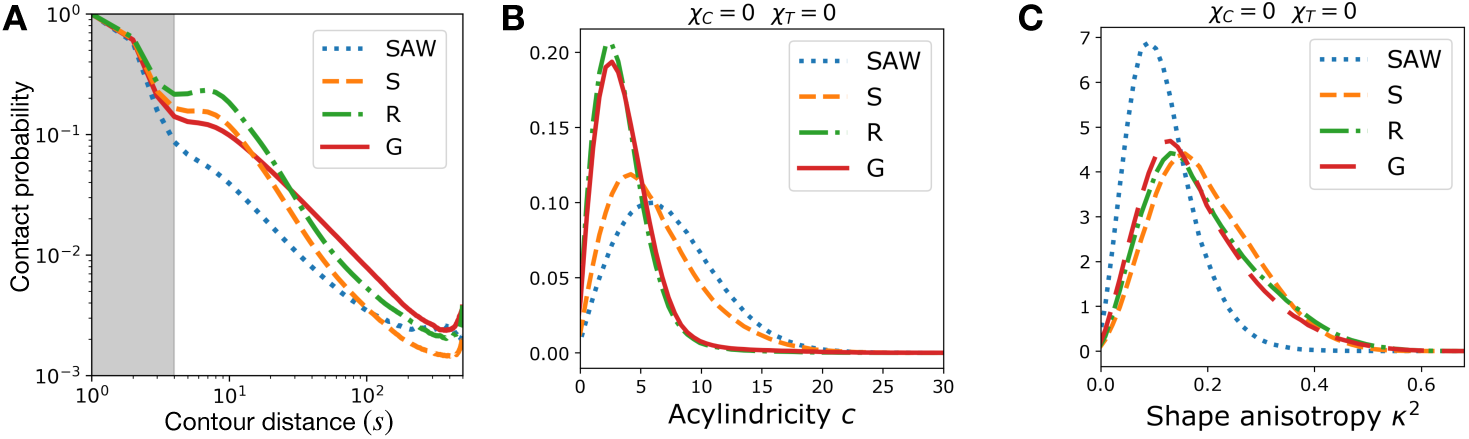
Lengthwise compaction shapes chromosomes. (A) Contact probability of intra-chromosome segments as a function of contour distance. The contact probability of a monomer with its neighbors, up to 3 or 4 monomers are mainly controlled by the angle-restrain part of of the homopolymer potential. The contact probability between the monomers beyond about 5 monomers is controlled by lengthwise compaction. (B) Acylindricity and (C) Shape anisotropy calculated from the Gyration tensor for chromosomes with different lengthwise compaction intensity: SAW, Globular Rope, and String phenotypes. Higher lengthwise compaction introduces cylindrical asymmetry in chromosomes structure.

**FIG. S3.**
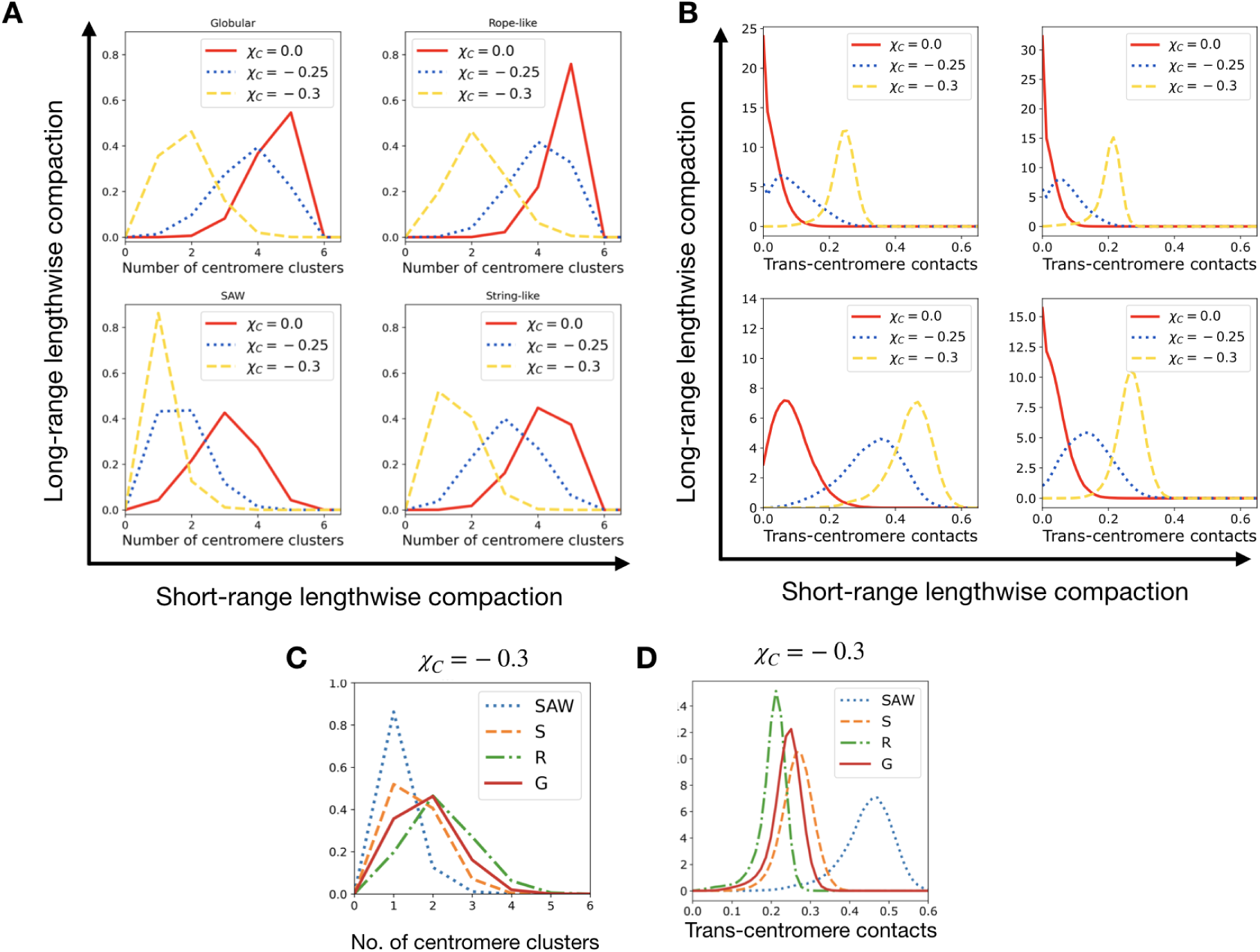
Centromere clustering is enhanced by centromere self-adhesion and counteracted by lengthwise compaction. (A) Number of centromere clusters and (B) Proportion of trans-centromere contacts, shown under different centromere self-adhesive interaction *χ*_*C*_, for various long- and short-range lengthwise compacted chromosomes. (C) Number of centromere clusters and (D) Trans-centromere contacts under *χ*_*C*_ = −0.3 for various lengthwise compaction.

**FIG. S4.**
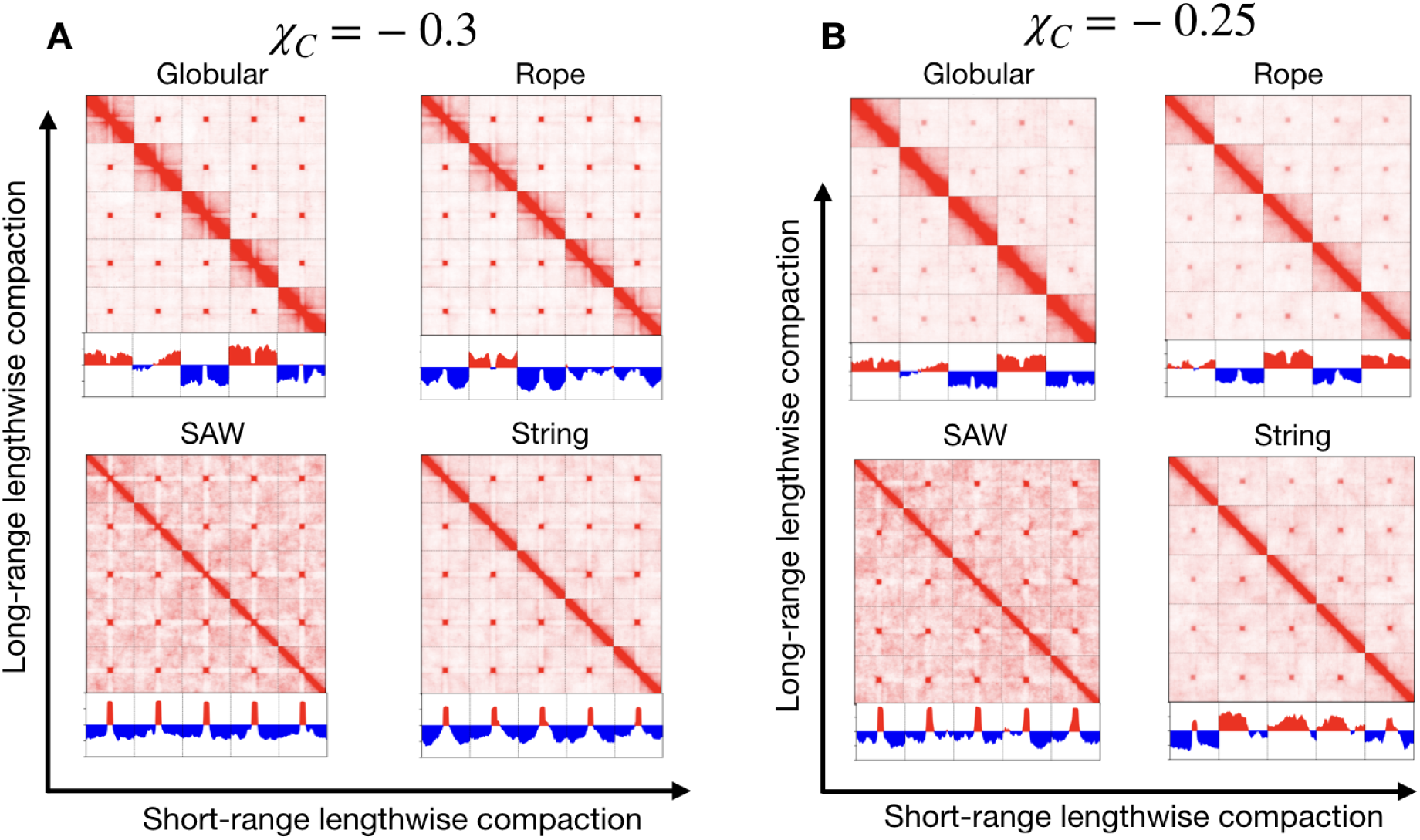
Contact matrices for different lengthwise compaction with centromere adhesion. Contact matrices and principal eigenvectors for SAW, G, R and S states with (A) strong (*χ*_*C*_ = −0.3) and (B) moderate (*χ*_*C*_ = −0.25) centromere adhesion.

**FIG. S5.**
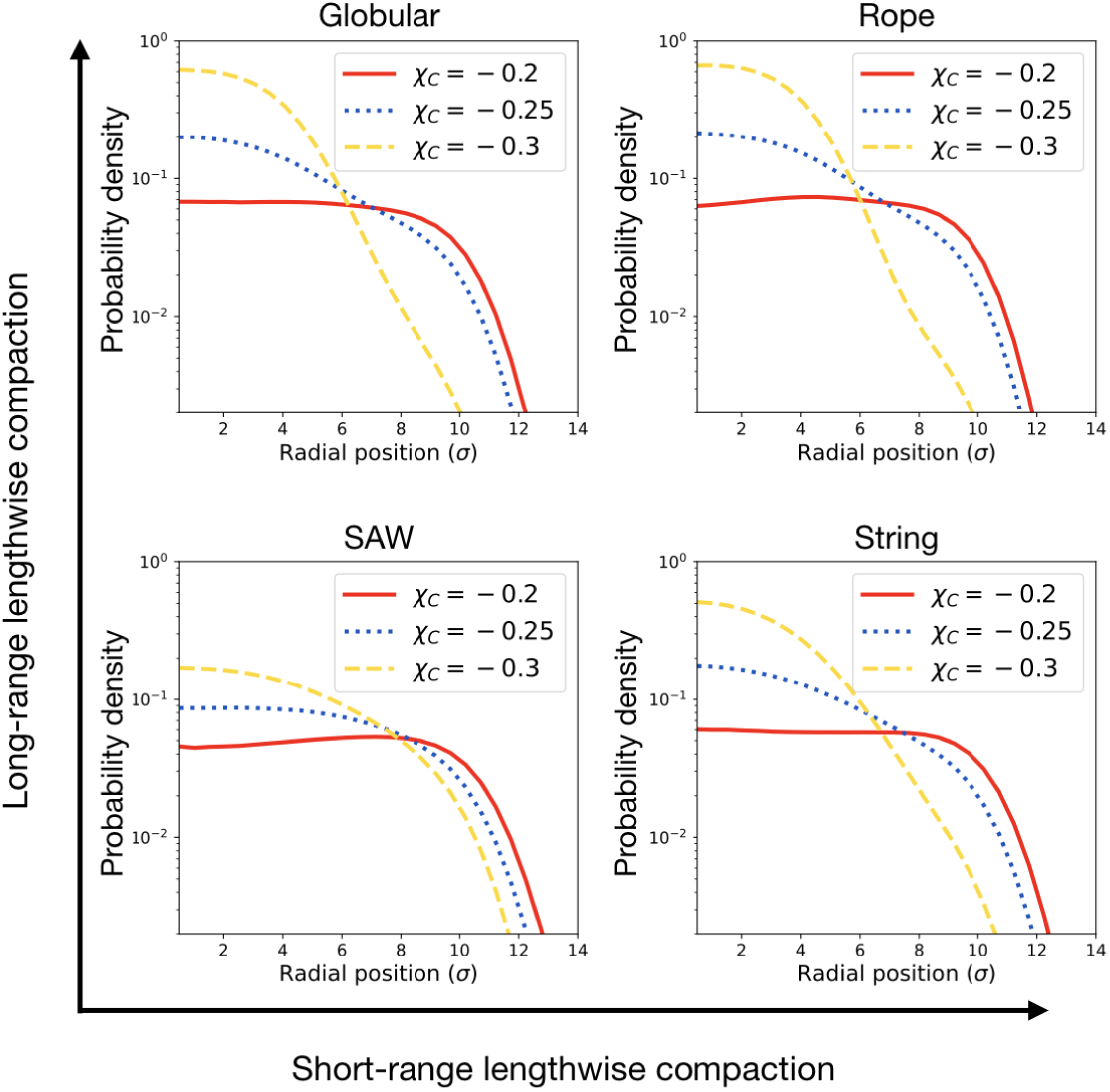
Compact centromeres with strong self-adhesion move to the center of the nucleus. Radial probability density of centromeres under various self-adhesion and lengthwise compaction shown. Note that the compact centromeres corresponding to Globular, Rope, and String-phenotypes when strongly adhere to other centromeres tend to localize near the center of the nucleus.

**FIG. S6.**
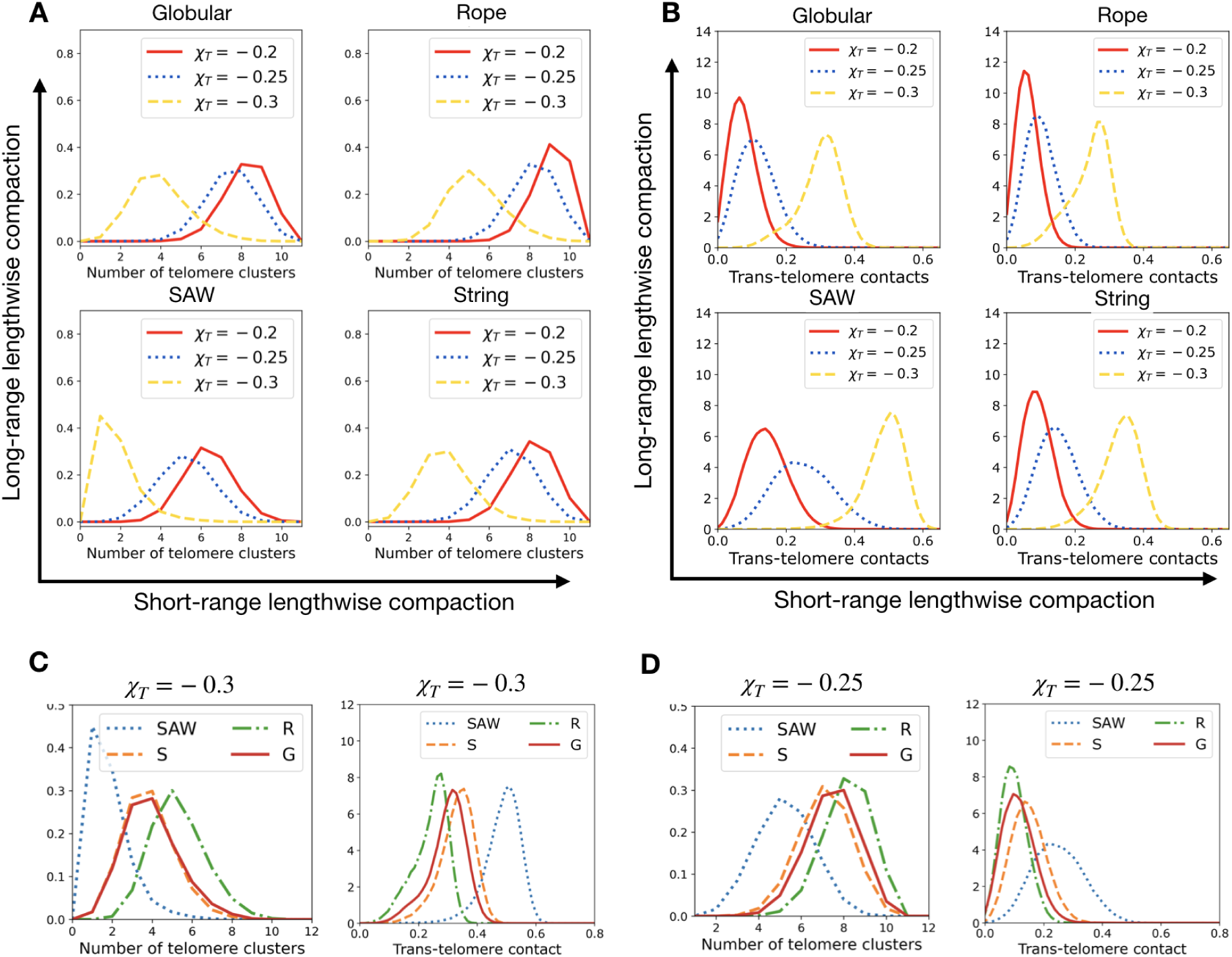
Telomere clustering is enhanced by telomere self-adhesion and counteracted by lengthwise compaction. (A) Number of telomere clusters (B) Proportion of trans-telomere contacts shown for various telomere adhesive interactions *χ*_*T*_ and under different lengthwise compaction. Number of telomere clusters and trans-telomere contacts for (C) *χ*_*T*_ = −0.3 and (D) *χ*_*T*_ = −0.25 compared for SAW, G, R and S-states.

**FIG. S7.**
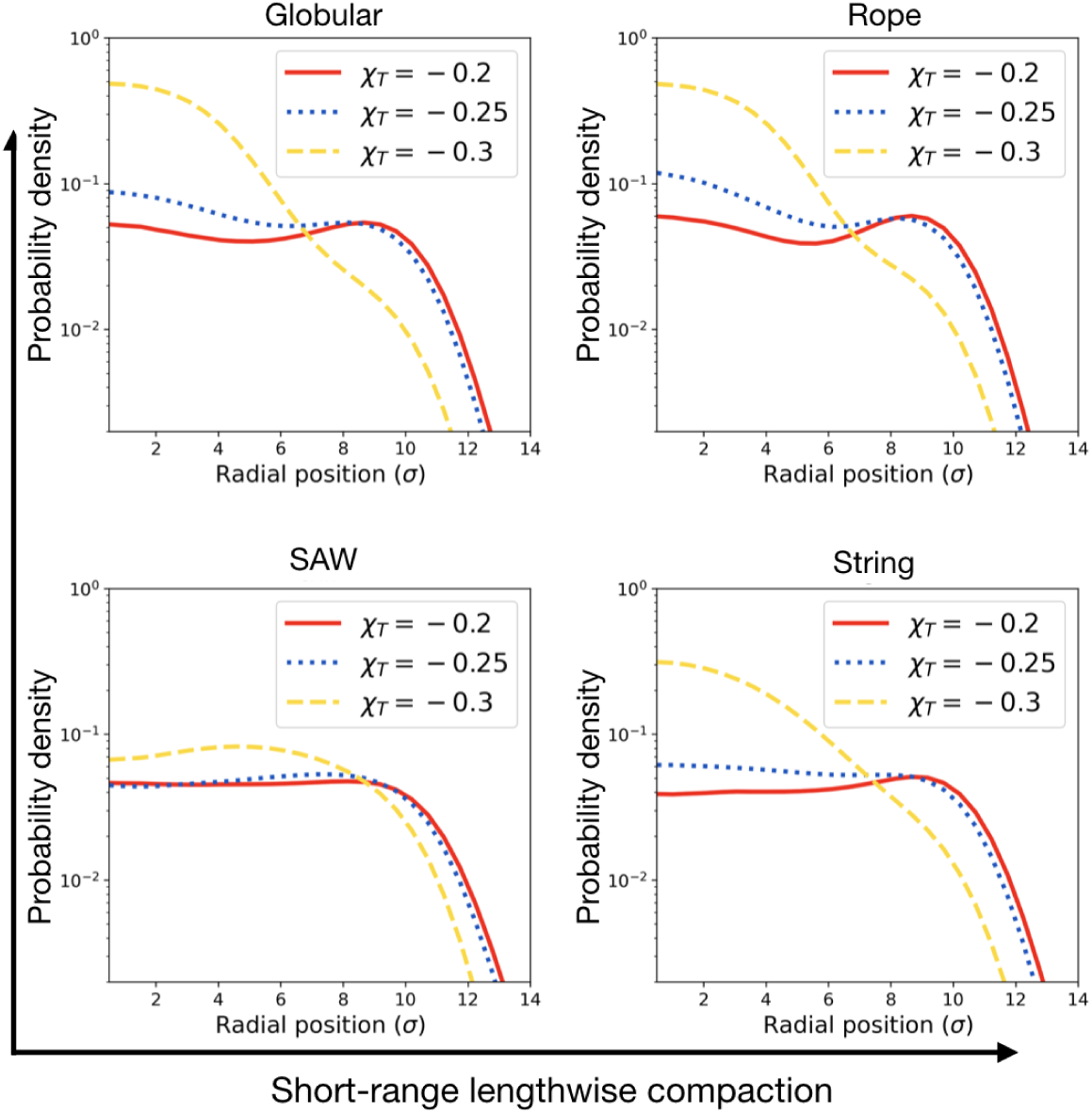
Radial density profile of telomeres. Radial probability density of telomeres under various self-adhesion and lengthwise compaction shown. Telomeres without preferential self-adhesion reside near the periphery when lengthwise compaction is high. This is likely due to the stiffness of the chromosome backbone that tends to place its ends near the periphery. However, upon strong self-adhesion, the telomeres move to the interior, much like the centromeres under similar conditions.

**FIG. S8.**
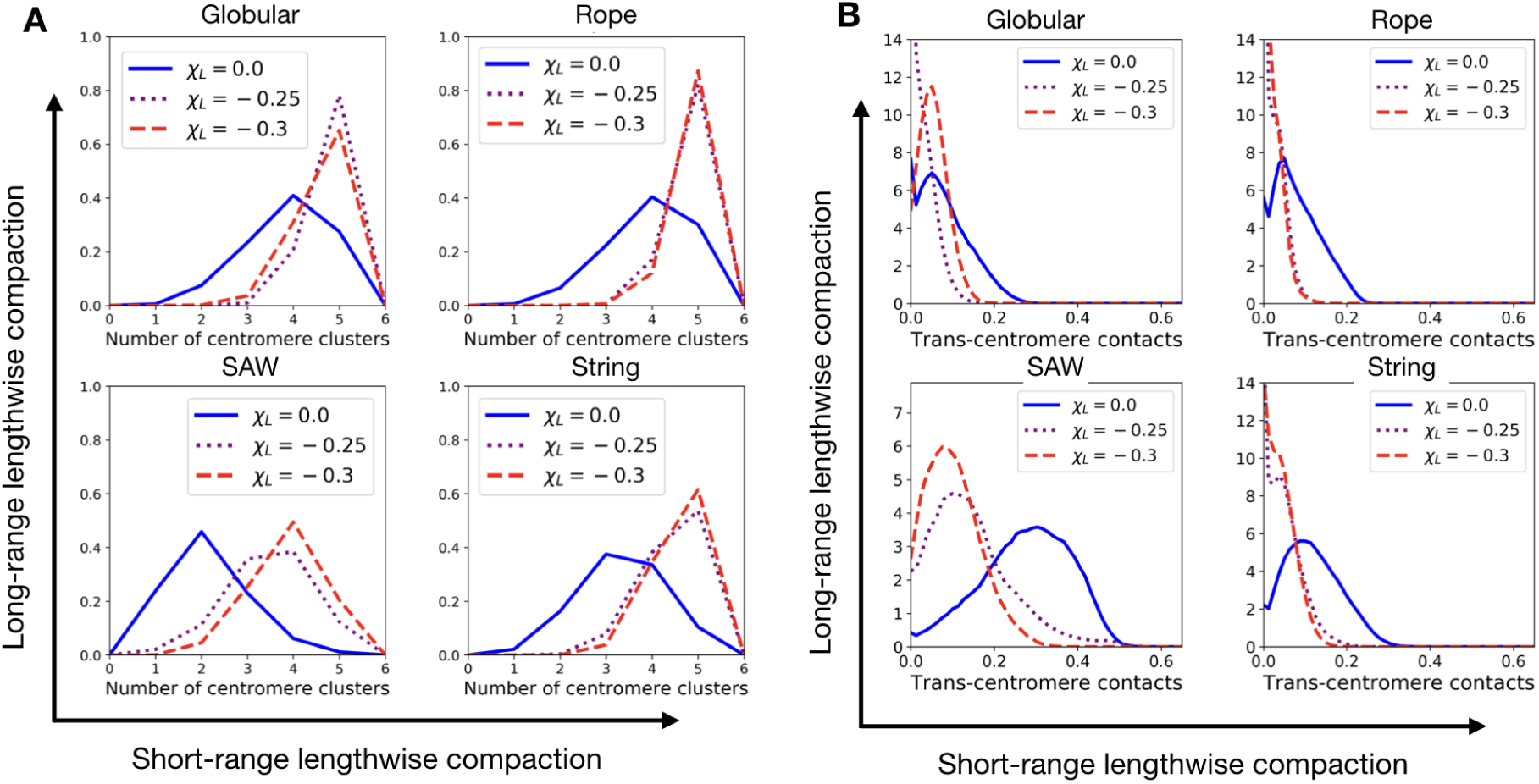
Lamina adhesion of centromeres counteract their clustering. (A) Number of centromeres and (B) Proportion of trans-centromere contacts for moderate (*χ*_*C*_ = −0.25) centromere self-adhesion but various lamina-tethering intensity *χ*_*L*_, showing higher *χ*_*L*_ increases the number of centromere clusters, and reduces the propensity of inter-centromere contacts.

**FIG. S9.**
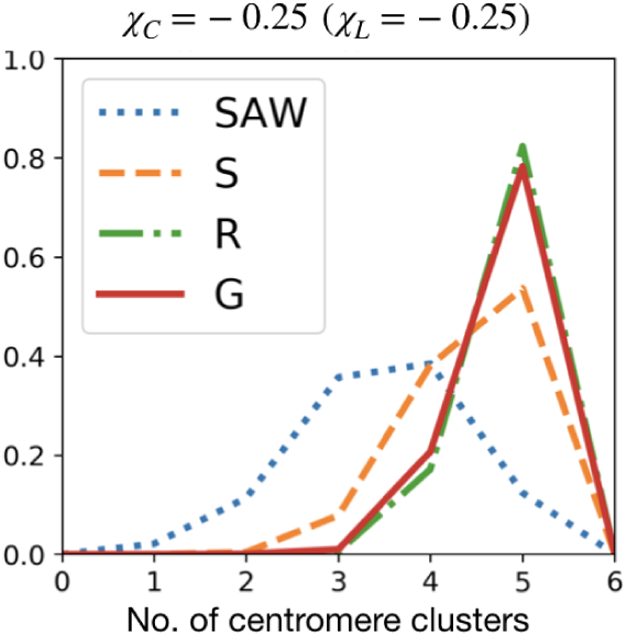
Lengthwise compaction counteracts centromere clustering in presence of lamina adheison. Number of centromere clusters for chromosomes with moderate lamina adhesion of centromeres (*χ*_*L*_ = −0.25) and centromere self adhesion (*χ*_*C*_ = −0.25) under various lengthwise compaction.

**FIG. S10.**
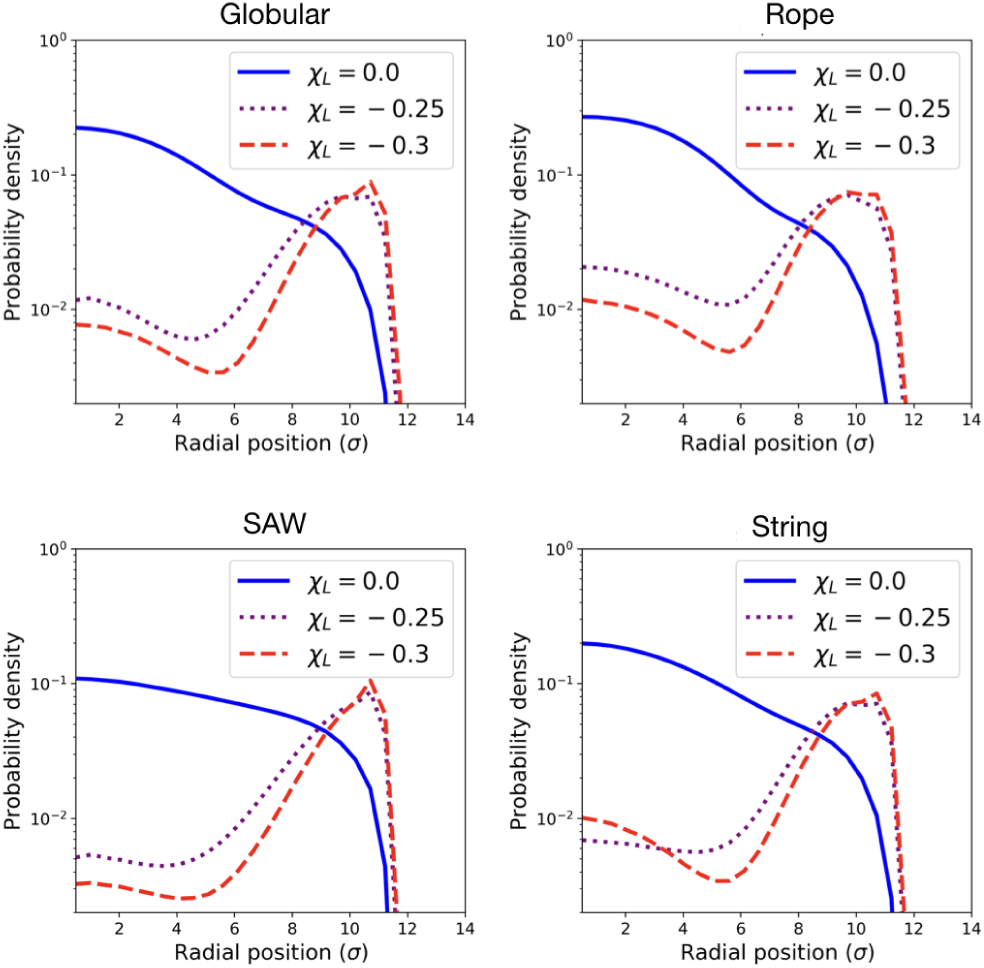
Radial density profile of centromeres with lamina adhesion. Radial density profile is shown for centromeres with self-adhesive intensity *χ*_*C*_ = −0.25. Centromeres tend to move to the periphery when interacting favorably with the lamina.

**FIG. S11.**
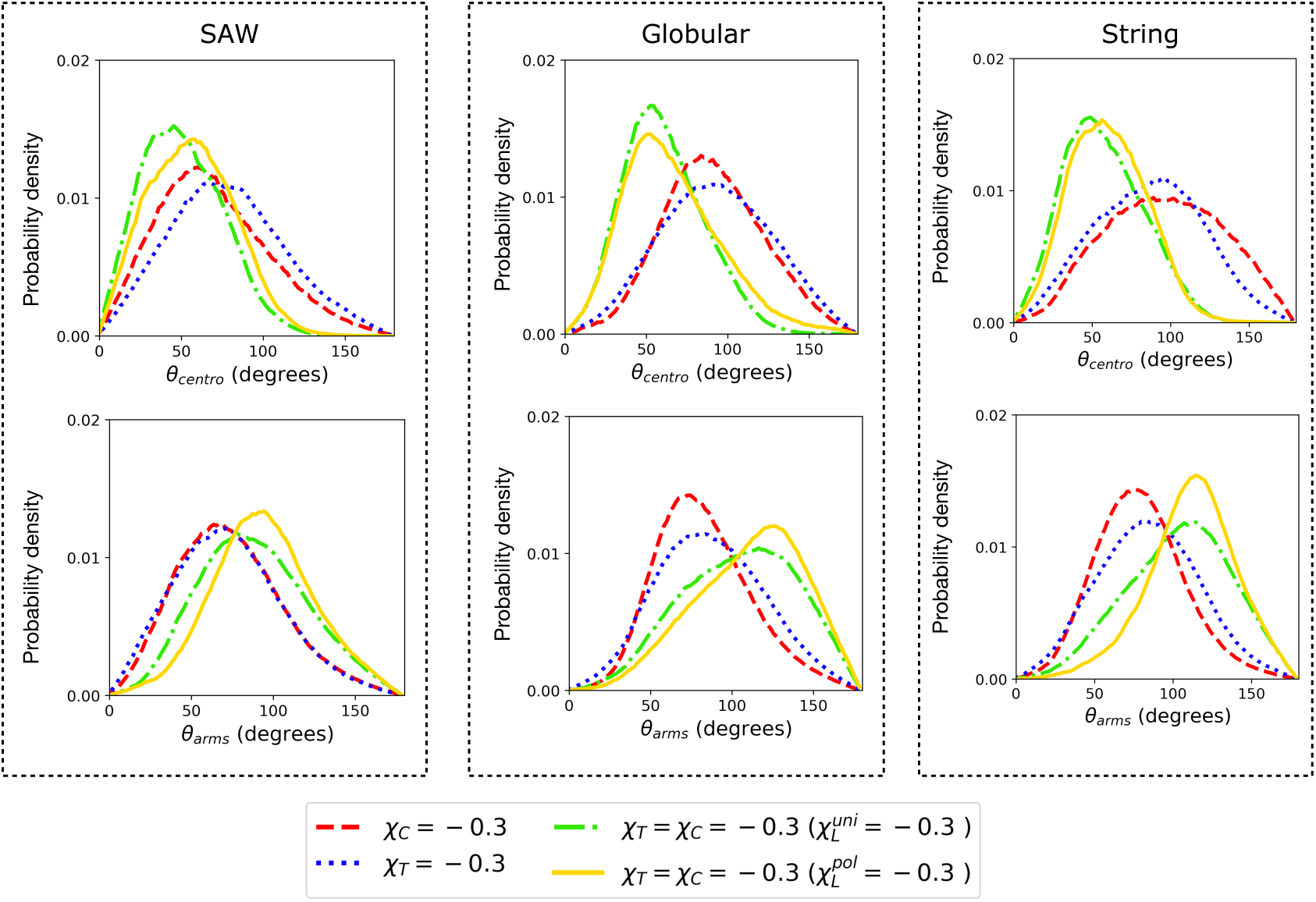
In presence of strong lamina tethering of centromeres and telomeres, length-wise compaction may aid fold over. Fold over angles: *θ*_*centro*_ and *θ*_*arm*_ for various cases, legend is the same as the Fig. 4D,E. The three panels correspond to different lengthwise compaction states, the rope-like phenotype is shown in Fig. 4.

**FIG. S12.**
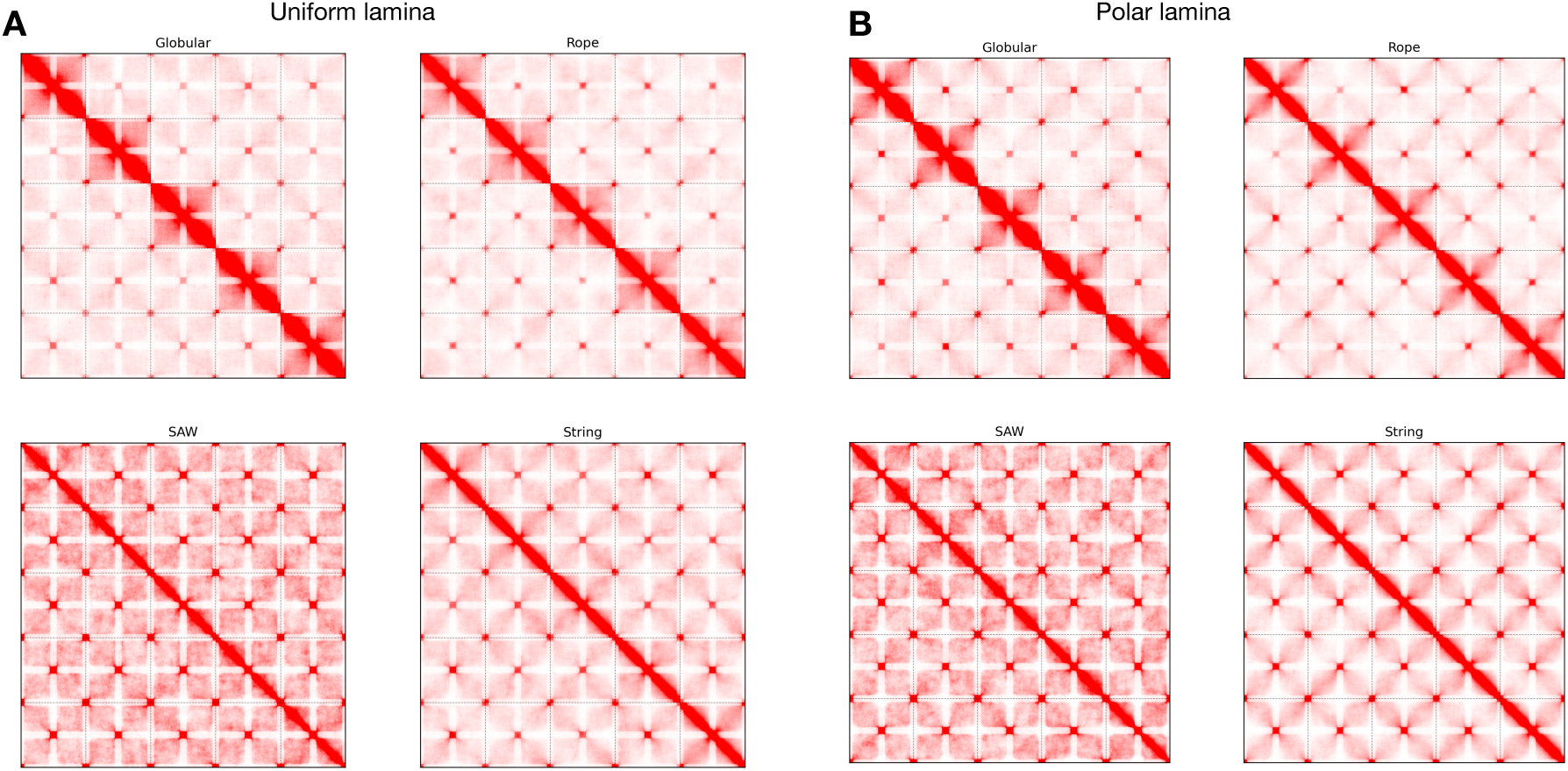
Contact maps showing fold over for polar and uniform lamina interactions. The contact maps for (A) uniform and (B) polar lamina interactions of strnegth *χ*_*L*_ = −0.3 for various lengthwise compaction states. All the panels correspond to strong centromere and telomere adhesion (*χ*_*C*_ = *χ*_*T*_ = −0.3).

**FIG. S13.**
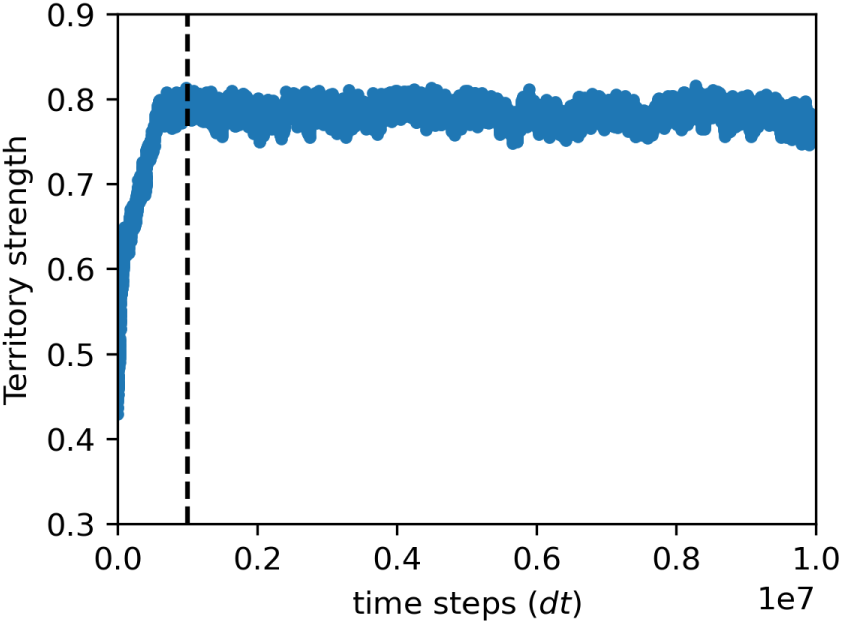
Steady state trajectories. The territory signal is shown for a single replica as a function of simulation time. The system was initialized as a SAW (hence the low territory at zero time), but simulated under the potential for G. The system reaches its steady state corresponding to G chromosomes in less than 10^6^ time steps. We exclude the initial one million time steps from our analysis (dashed line), to ensure that the memory of the initial configuration is completely lost, and we are capturing the steady state.

